# Structures of the Ndc80 complex and its interactions at the yeast kinetochore-microtubule interface

**DOI:** 10.1101/2022.11.09.515846

**Authors:** Jacob Zahm, Simon Jenni, Stephen Harrison

**Affiliations:** Department of Biological Chemistry and Molecular Pharmacology, Harvard Medical School, Boston, MA 02115, USA; Howard Hughes Medical Institute, Harvard Medical School, Boston, MA 02115, USA

## Abstract

The conserved Ndc80 kinetochore complex, Ndc80c, is the principal link between spindle microtubules and centromere associated proteins during chromosome segregation. We used AlphaFold 2 to obtain structural predictions of the Ndc80 “loop” region and the Ndc80:Nuf2 globular head domains that interact with the Dam1 subunit of the decameric DASH/Dam1 complex (Dam1c). The predictions guided design of constructs that readily yielded crystal structures, essentially congruent with the predicted ones. The Ndc80 “loop” is a stiff, straight α-helical “switchback” structure, and flexibility within the long Ndc80c rod occurs instead at a hinge point between the globular head and the loop. Conserved stretches of the Dam1 C terminus bind Ndc80c with a short α helix followed by an extended segment such that phosphorylation of Dam1 serines 257, 265, and 292 by the mitotic kinase Ipl1/Aurora B can release this contact during error correction of mis-attached kinetochores. We integrate the structural results presented here into our current molecular model of the kinetochore-microtubule interface. The model illustrates how multiple interactions between Ndc80c, DASH/Dam1c and the microtubule lattice stabilize kinetochore attachments.

## INTRODUCTION

Kinetochores, essential components of the chromosome segregation machinery, connect centromeres to the mitotic spindle. The bridge between centromere-associated kinetochore assemblies and spindle microtubules is the Ndc80 complex (Ndc80c), a heterotetramer, conserved from yeast to mammals, with globular elements at both ends of a ~600 Å long, largely α-helical coiled-coil shaft (Fig. 1) (Ciferri et al., 2005; Wang et al., 2008; Wei et al., 2005). Closely paired, calponin homology (CH) domains of Ndc80 and Nuf2 at one end, augmented by a flexible, N-terminal extension of Ndc80, form a microtubule-attachment module (Alushin et al., 2010; Ciferri et al., 2008; Wei et al., 2007). Tightly associated, RWD domains of Spc24 and Spc25 at the other end bind adaptor proteins emanating from centromere-bound, kinetochore components (Dimitrova et al., 2016; Malvezzi et al., 2013; Nishino et al., 2013; Wei et al., 2006). The coiled-coil shaft of Ndc80:Nuf2 has two specializations -- a kink or hinge at about 160 Å from the globular end (Wang et al., 2008) and a so-called “loop” at about 240 Å. The latter structure is effectively an insertion of ~50–80 residues into Ndc80 opposite a continuous helix in Nuf2 (Maiolica et al., 2007). The shafts of the two heterodimeric components -- Ndc80:Nuf2 and Spc24:Scp25 -- meet in an overlapping, four-way junction, or tetramerization domain, included in the crystal structure of a shortened, “dwarf” Ndc80c (Valverde et al., 2016).

**Fig. 1.**
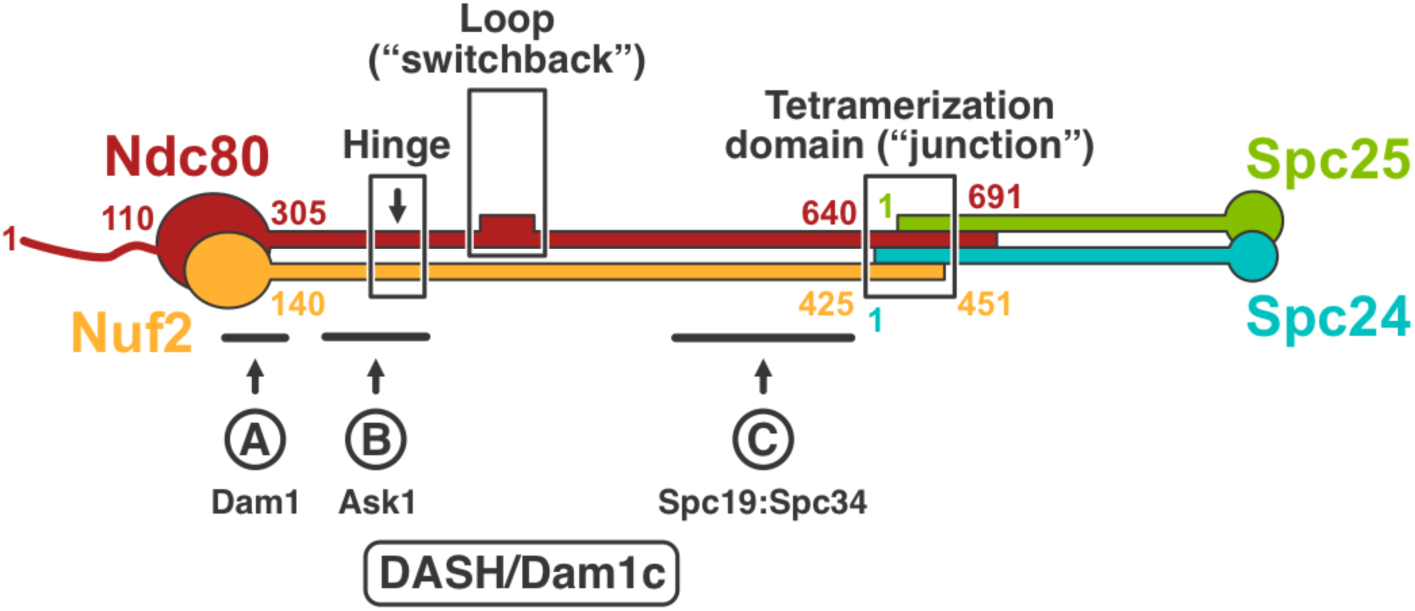
Schematic diagram of Ndc80c. Residue numbers for key junction points in the *S. cerevisiae* sequences shown. Solid ovals represent the calponin homolog (CH) domains of Ndc80 (red) and Nuf2 (orange) at one end and the RWD domains of Spc24 (blue) and Spc25 (green) at the other. The positions of the Ndc80 hinge, loop, and the four-way junction are boxed and labeled, respectively. The locations of the three interactions, defined by crosslinking mass-spectrometry (Kim et al., 2017), between Ndc80c and C-terminal extensions of DASH/Dam1c subunits are indicated. Interaction A involves Dam1, interaction B, Ask1, and interactions C, Spc19:Spc34.

In yeast, several additional kinetochore components bind at various positions along the Ndc80c. At least three interactions involve subunits of the DASH/Dam1-complex (Dam1c, consisting of Ask1, Dad1, Dad2, Dad3, Dad4, Dam1, Duo1, Hsk3, Spc19, and Spc34) (Jenni and Harrison, 2018; Miranda et al., 2005; Westermann et al., 2005), which coordinates end-on attachment of multiple Ndc80c rods around the splayed plus-end of a kinetochore microtubule (Lampert et al., 2013). A fourth contacting component is the microtubule polymerase, Stu2 (Miller et al., 2016). Fig. 1 shows positions of the DASH/Dam1c interactions, as mapped by genetic and biochemical studies including mass spectrometric analysis of cross-linked peptides (Flores et al., 2022; Kim et al., 2017; Lampert et al., 2013). The Dam1 subunit binds at the base of the Ndc80:Nuf2 globular “head”, Ask1 along the shaft between the globular head and the hinge, and Spc19:Spc34 between the loop and the four-way junction, respectively. All these interactions between Ndc80c and DASH/Dam1c are regulated by phosphorylation of certain serine or threonine residues (Cheeseman et al., 2002; Flores et al., 2022). The Stu2 interaction, visualized by x-ray crystallography, is with the four-way junction (Zahm et al., 2021).

Budding yeast kinetochores first attach to the lateral surface of spindle microtubules (Tanaka et al., 2005). This initial capture must convert to end-on attachment to produce a structure that can track with microtubule shortening and bear the load thereby generated (Grishchuk et al., 2008; Umbreit et al., 2014). The conversion occurs when the end of a disassembling microtubule reaches the laterally attached kinetochore (Kitamura et al., 2007; Tanaka et al., 2007; Tanaka et al., 2005). The DASH/Dam1c ring is dispensable for lateral capture but essential for conversion to end-on capture (Tanaka et al., 2007). Deletions and certain mutations in the Ndc80 loop block this conversion (Maure et al., 2011). The microtubule polymerase, Stu2, also appears to have a role (Miller et al., 2016; Tanaka et al., 2007). As steps towards working out the molecular events in conversion from lateral to end-on attachment, we have extended our previous studies of the Stu2 interaction at the Ndc80c four-way junction (Zahm et al., 2021) by determining crystal structures of the Ndc80 loop and of the Dam1 association at the base of the Ndc80:Nuf2 globular head. The latter structure explains why phosphorylation of Dam1 serine residues 257, 265, and 292 by Ipl1/AuroraB inhibits attachment (Cheeseman et al., 2002; Jin et al., 2017; Keating et al., 2009). We compare the new structures with the AlphaFold 2 (AF2) (Jumper et al., 2021) predictions that motivated design of the constructs we used, and we show that the AF2 prediction of the Ndc80c’s hinge point agrees with the position of selective proteolytic sensitivity described in our early studies of Ndc80c architecture (Wei et al., 2005).

## RESULTS

### Ndc80 loop

Ndc80 amino-acid sequences across the region that contains the loop align well for a set of budding yeasts (Fig. 2A) and also for a representative set of metazoans (Fig. 2B), but the sequences align poorly between those two groups (Data S1–S4). Structure predictions from multi-chain AF2 for segments of Ndc80:Nuf2 that include the loop and flanking sequences are thus, as expected, very similar across the budding yeasts and across metazoans, and overall quite similar between those two groups, but distinct in some local features (Fig. 2C). An essentially identical prediction has motivated recent functional studies for the human Ndc80 loop by another group, as discussed at the end of this paper (Polley et al., 2022). Both sequences and predicted structures for fission yeast (*Schizosaccharomyces pombe*) and fungi in phyla other than Saccharomycetes align, in predicted structure and in sequence, with the metazoans (Figs. 2B and C). We expressed segments of both the yeast and human Ndc80 polypeptide chains and the respective paired segments (from the prediction) of Nuf2. We obtained crystals of the human complex in space group C_2_ (a = 49.9 Å, b = 72.3 Å, c = 80.1 Å, β = 98.56°). From merged datasets that extended to a minimum Bragg spacing of 2.0 Å, we determined the structure by molecular replacement with a four-part model based on the AF2 prediction (see Materials and Methods for details and for controls to eliminate model bias, Table S1 for data and refinement statistics, and Fig. S1 for representative density maps). The final model (Fig. 2D) agrees well with the predicted structure (Fig. S2), consistent with the strength and robustness of the molecular replacement calculations.

**Fig. 2.**
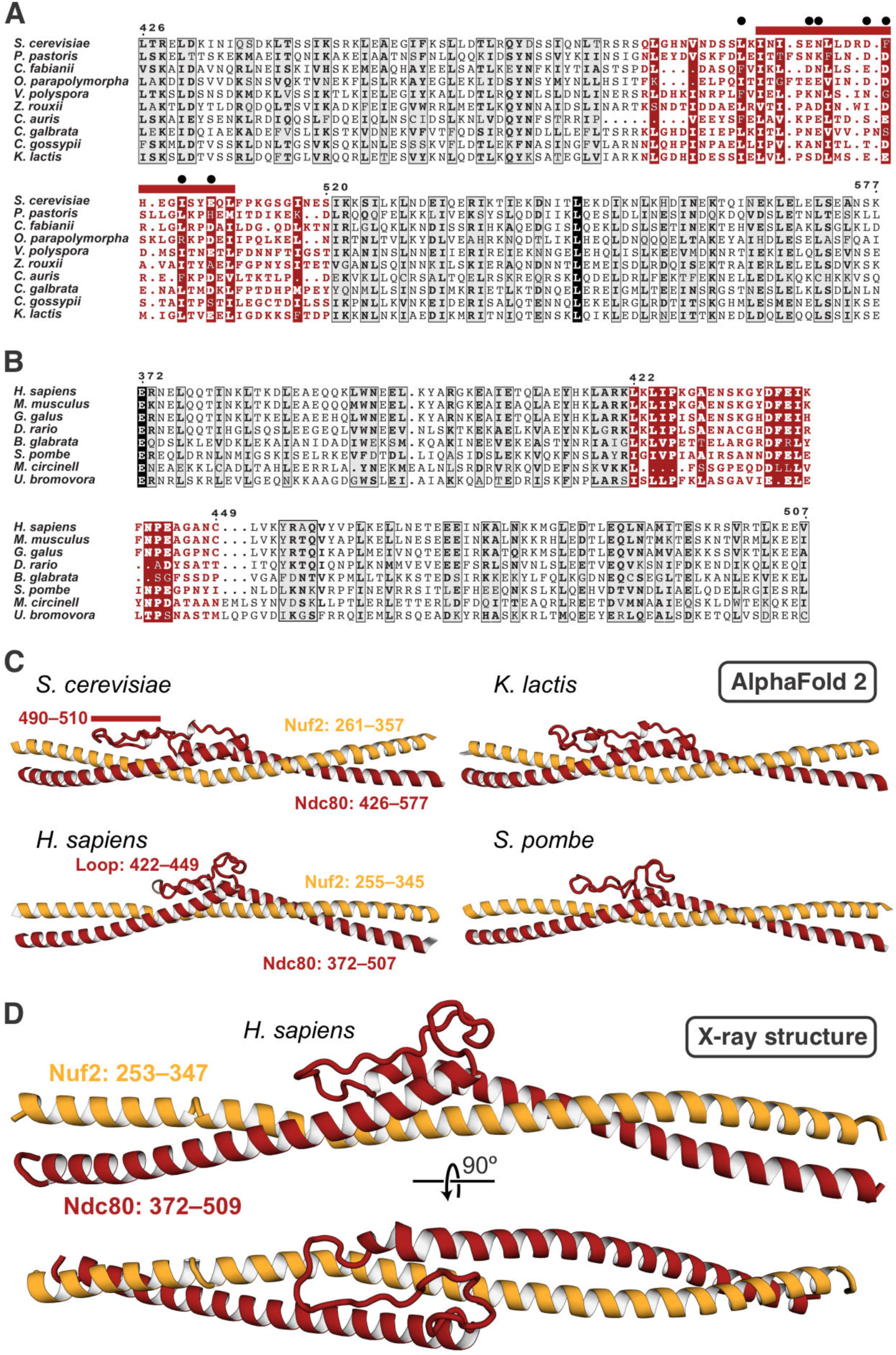
The Ndc80 loop. (A) Sequence alignment for the Ndc80 loop and adjacent residues for budding yeasts. Numbers above the sequences are for *S. cerevisiae*, residues 426–577). The loop residues are in red. The red bar above the alignment corresponds to the longest segment that can be deleted without lethality (Maure et al., 2011). The black dots are residues mutated to alanine in the temperature-sensitive mutant *ndc80-7A* (Maure et al., 2011). (B) Sequence alignment for the Ndc80 loop and adjacent residues for metazoans plus non-budding fungi. Numbers correspond to *H. sapiens*, residues 372–507. The loop residues are in red. (C) AF2 predictions for two budding yeasts, human, and fission yeast. The extended loop that doubles back between the two overlapped helices at either end is substantially longer in budding yeast, with the structural difference coinciding almost precisely with the longest non-lethal deletion (red bar, residues 490–510). For illustrations of the predictions with AF2 confidence levels, see Fig. S2A. (D) Cartoon representation of the crystal structure of the human loop segment of Ndc80:Nuf2. The view of the upper cartoon is the same as the view of the prediction for the human structure in panel (C). A full comparison is in Fig. S2.

Both the structure and the AF2 prediction show that the “loop” is in fact an α-helical “switchback”, in which the helix entering the loop from the N-terminal side continues beyond its pairing with Nuf2, reverses direction into a segment with a partly disordered loop (using “loop” in the stricter sense) as it turns back into an α helix, which continues into the region paired again with Nuf2. The Nuf2 chain is a continuous helix opposite the entire switchback. The short, disordered segment in the human Ndc80 structure corresponds (in the predictions) to the segment of *Saccharomyces cerevisiae* Ndc80 containing the substitutions in a seven-alanine mutant shown to fail in the transition to end-on attachment in budding yeast (Maure et al., 2011) (Fig. 2A). The very close correspondence of predicted and experimental loop structures for human Ndc80:Nuf2 give us confidence in the predictions for other species, including the budding yeasts. The success of predictions for the head-shaft transition and its interaction with Dam1 described below also supports the credibility of the AF2 models.

### Ndc80:Nuf2:Dam1 interaction

Genetic, biochemical, and crosslinking data suggest that a segment of the extended C-terminus of the Dam1 subunit binds with a region of Ndc80:Nuf2 near the position at which the Ndc80 and Nuf2 polypeptide chains transition from their paired, CH domains into a helical coiled-coil (Flores et al., 2022; Kim et al., 2017; Lampert et al., 2013). Various trial calculations in multi-chain AF2, in which we varied the details of the Dam1 segment we included, led to a self-consistent prediction for binding, in which two stretches of conserved Dam1 residues (252–270 and 290–305) (Fig. 3A and Data S5) showed the highest confidence scores. The predicted structure of the docked segment suggested that a fusion construct of part of the Dam1 C terminus (residues 252–305) with the Nuf2 N terminus, combined with a deletion of residues between the two conserved stretches (residues 271–289), might ensure high local concentration and increase the likelihood of obtaining informative crystals. We therefore made the inferred Ndc80:Dam1-Nuf2 fusion construct as part of the dwarf Ndc80c, expressed and purified the recombinant protein, and obtained crystals in space group P3_2_12 (a = b = 130.0 Å, c = 216.3 Å). Molecular replacement with the Ndc80:Nuf2 component from PDB-ID 5TCS (Valverde et al., 2016) and initial refinement without any Dam1 model yielded unbiased difference density for the Dam1 segment (Fig. S3), allowing us to build and refine a model (data and refinement statistics in Table S1). The Spc24:Scp25 component had dissociated during crystallization, and only the Ndc80:Dam1-Nuf2 heterodimer was present in the crystals. The model and a comparison with the AF2 prediction are in Figs. 3B and C. The regions of the prediction with good confidence scores correspond well to the experimental result (Fig. S4). A small difference in the short α helix (residues 258–267) does not substantially reorganize any of the side-chain interactions.

**Fig. 3.**
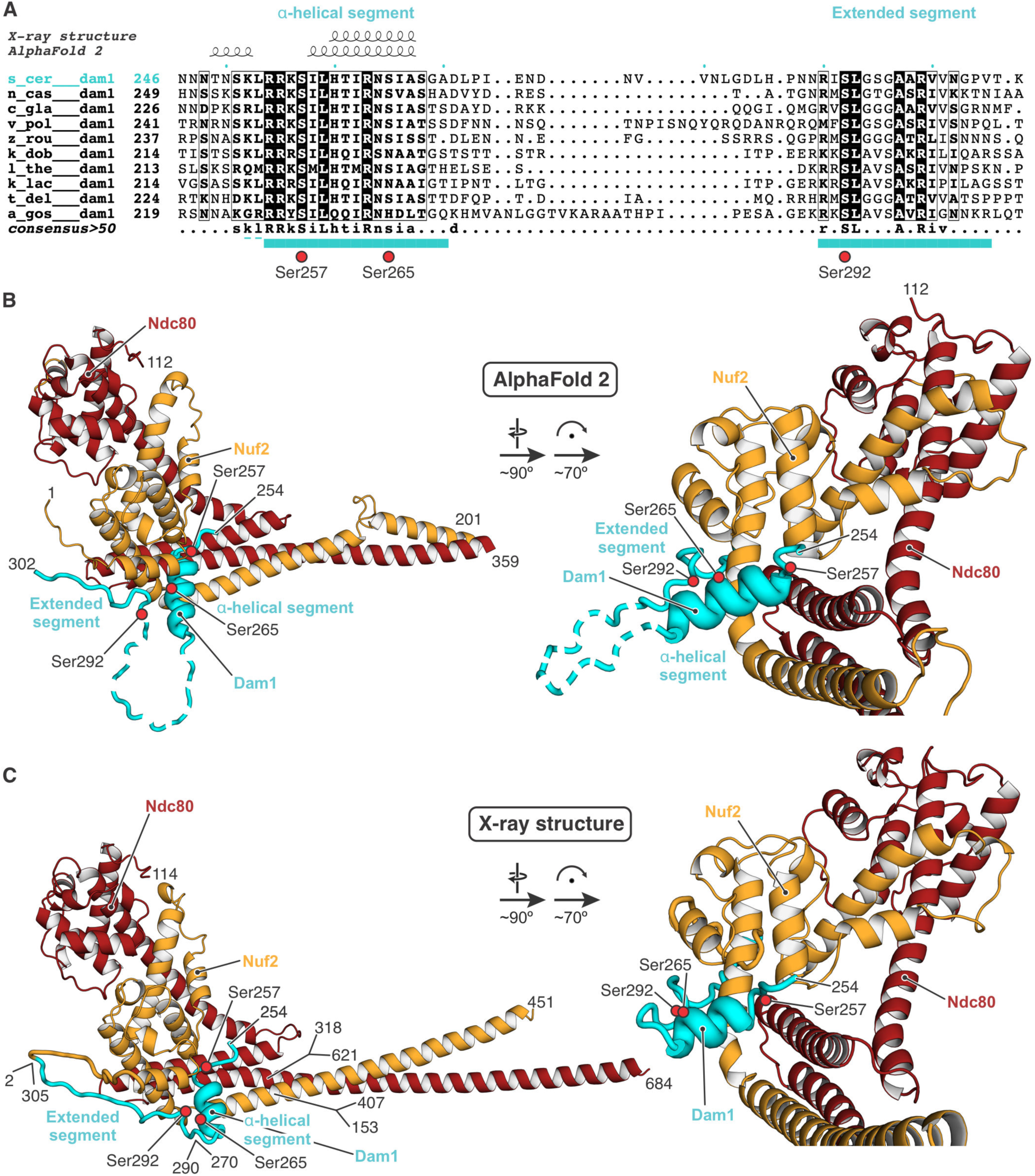
Association of a yeast Dam1 segment with the globular “head” of Ndc80:Nuf2. (A) Sequence alignment for budding yeasts, showing the Dam1 region of *S. cerevisiae* residues 246– 307 (the full alignment is shown in Data S3). Cyan bars below the sequences are residues of the conserved α-helical and extended segments, respectively, that were modeled in the crystal structure here. Dashed lines are residues that were included in the construct but not modeled. Serine residues 257, 265, and 292, which can be phosphorylated by Ipl1, are indicated by red circles. (B) AF2 prediction for the residues shown. The dashed loop in Dam1 had very low reported confidence values and predicted variably from run to run. (C) Cartoon representation of the crystal structure of the Ndc80:Dam1-Nuf2 fusion, residues as shown. The discontinuities in Ndc80 and Nuf2 are due to use of the Ndc80c “dwarf” design to produce the construct (Valverde et al., 2016). The deletion of 19 residues in Dam1 corresponds to the low-confidence loop in (B). The “prediction” for the globular domains is essentially precise, as the coordinates for the same sequences are in the PDB. Note that the position of the short Dam1 α helix is shifted by about 3 Å from its position in the prediction (Fig. S4C and E). Red circles are the positions of serines 257, 265 and 290 (OG oxygen atoms). See also Fig. S4.

Dam1 binds Ndc80:Nuf2 with an α-helical segment, which rests in a cleft between the coiled coil and the globular head domains, and an extended segment, which runs along the N terminus of Nuf2, but also contacts Ndc80 (Fig. 3B and C). The residues connecting the two segments are not conserved (Fig. 3A), predicted to be flexible by AF2, and were omitted in the construct of the crystal structure. Molecular details of the interactions are shown in Fig. 4. A salt bridge between Dam1 Arg299 and Ndc80 Asp295, both conserved across budding yeasts (Data S3 and S5), anchors the C-terminal extended segment of the Dam1 fragment (Fig. 4B). The Dam1 sequence in the structure also includes three serines (residues 257, 265, and 292) (Fig. 3A), the first and third conserved among budding yeasts and the second largely so, that are potential Ipl1 targets for regulating Ndc80c binding (Data S5). The structure suggests that phosphorylation of serines 257 and 265 would substantially weaken the contact (Fig. 4A). Ser257 is opposite conserved Glu276 of Ndc80 and its phosphorylation would lead to electrostatic repulsion; serine 292 is well exposed in the model, but the thermal parameter in that segment of polypeptide chain is relatively high (Fig. S4B), and electrostatic repulsion of a phosphoserine side chain by nearby negative charge might in principle weaken the contact and make phosphorylation of the other two sites more likely.

**Fig. 4.**
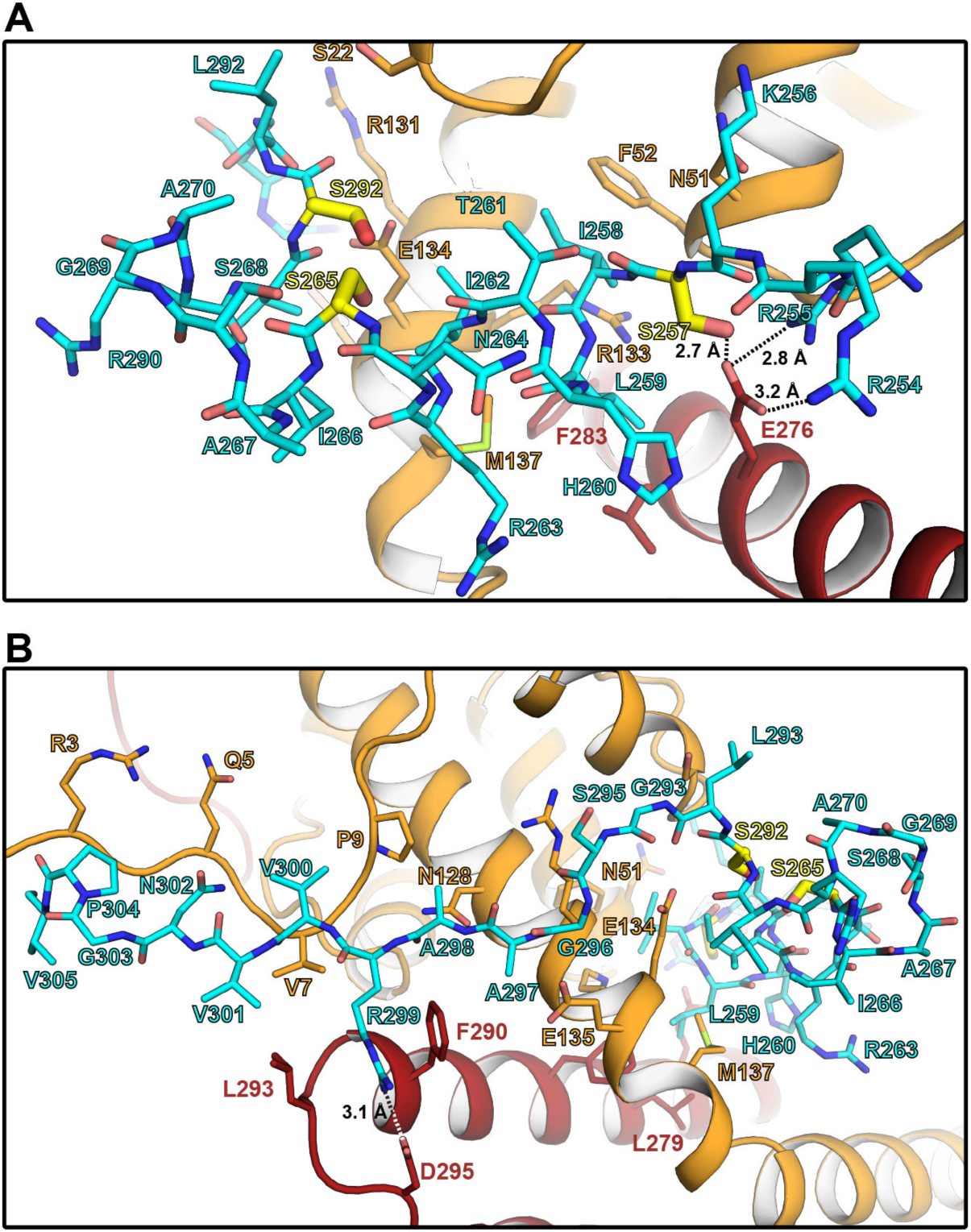
Close up of the Dam1:Ndc80c contacts. (A) Interactions between the Dam1 α-helical segment (cyan) and Ndc80:Nuf2 (red and orange, respectively). Serine residues 257, 265 and 292 that are targets for phosphorylation by Ipl1/Aurora B are shown in yellow. The side chain of Ser257 is close to conserved Ndc80 glutamate 276 and its phosphorylation would result in electrostatic repulsion. (B) Interactions between the Dam1 extended segment (cyan) and Ndc80:Nuf2 (red and orange, respectively). Note the salt bridge between conserved Dam1 residue Arg299 and Ndc80 Asp295.

### Hinge

Electron micrographs of intact, recombinant Ndc80 suggest the presence of a variable bend in the shaft between the Ndc80:Nuf2 globular domains and the four-way junction with Spc24:Spc25 (Huis In ’t Veld et al., 2016; Wang et al., 2008). Although the bend was initially proposed to coincide with the position of the loop, which was an evident discontinuity in the alignment of predicted Ndc80 and Nuf2 coiled-coil sequences, measurements of its average position in published electron micrographs suggested a location between the globular domains and the loop (Jenni et al., 2017), probably coincident with preferential proteolytic cleavage points, detected in our early work on the yeast Ndc80c, C-terminal to Ndc80 lysines 380 and 409 (Data S3) and Nuf2 lysines 220 and leucine 237 (Data S4) (Wei et al., 2005). AF2 predictions for Ndc80:Nuf2 across this region show a bend at Ndc80 residue 412 and Nuf2 residue 245, with some variation in the bend angle and direction depending on the length of the Ndc80:Nuf2 segment submitted for prediction. One of the cleavages is consistent with the predicted hinge position, a likely point of proteolytic sensitivity; the other is consistent with some instability of the exposed C-terminus of the coiled coil, allowing the protease(s) to cleave somewhat further toward the globular head. The position of the predicted hinge is also consistent with the observed, variable kink in electron micrographs (Ciferri et al., 2005). Fig. 5 incorporates this predicted hinge into an overall composite model of full-length Ndc80c.

**Fig. 5.**
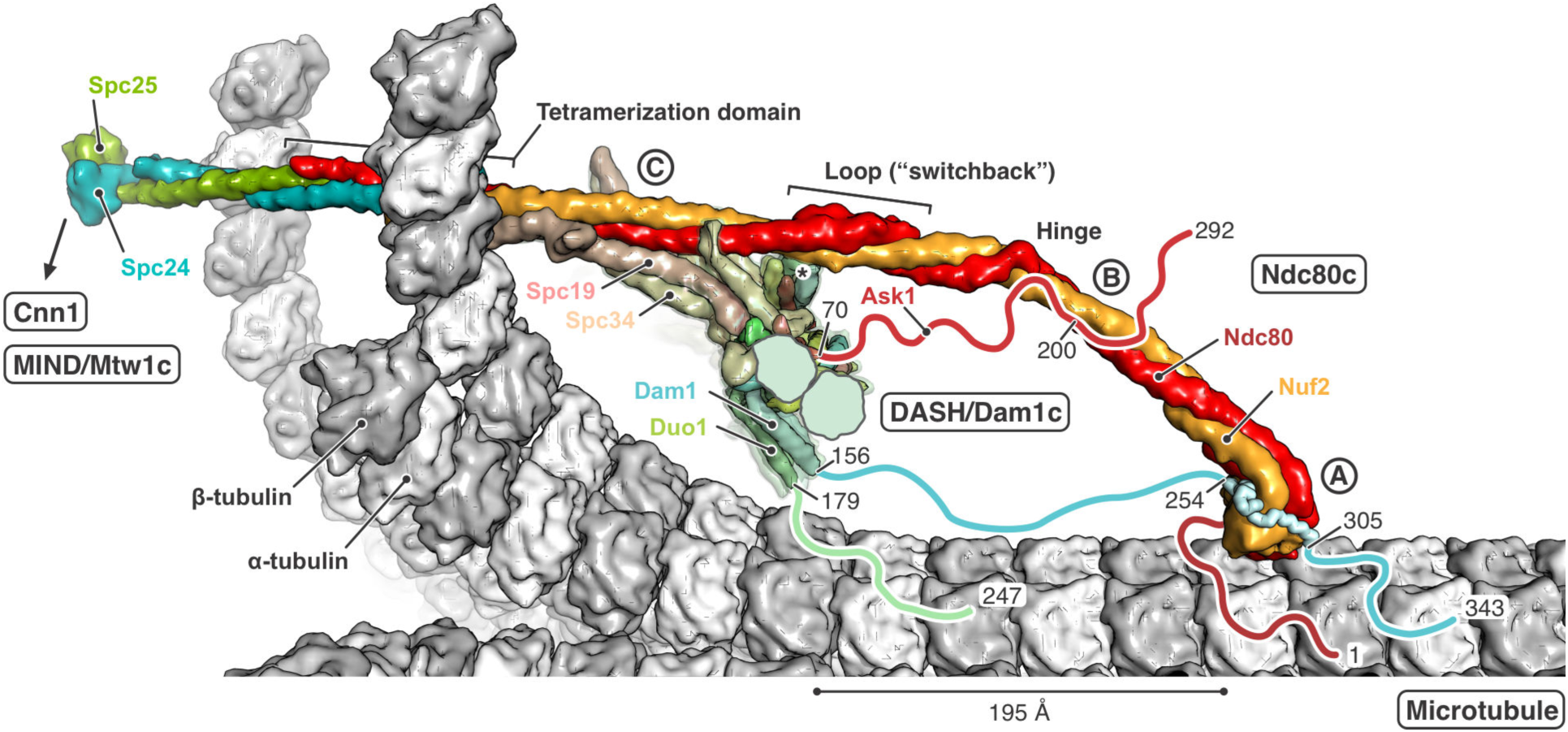
Model of the interactions of yeast Ndc80c with DASH/Dam1c and a kinetochore microtubule (MT). In the close-up view, a single Ndc80c in “end-on” attachment to a curled MT is shown. α- and β-tubulin subunits are colored white and gray, respectively. Ndc80c subunits are colored as in Fig. 1. The DASH/Dam1c ring, composed of 17 heterodecameric complexes, is partially cut for illustration. Flexible extensions for which interactions with their binding partner are not yet structurally characterized were omitted from the models and drawn schematically as lines, including the N terminus of Ndc80 (red) and the C termini of Ask1 (red), Dam1 (cyan), and Duo1 (green). The N termini of Dam1 and Duo1 are not shown; an asterisk indicates their location next to the Ndc80c loop. Relevant residues are numbered. The three interactions between Ndc80c and DASH/Dam1c are labeled A, B, and C (Jenni and Harrison, 2018; Kim et al., 2017). Interaction A is defined by binding of the C-terminal Dam1 extension to the Ndc80:Nuf2 head domains. The segment observed in the crystal structure here is shown in ribbon representation. Interaction B is between the C terminus of Ask1 and Ndc80:Nuf2 coiled coil residues before the hinge. Interaction C involves binding of the Spc19:Spc34 protrusion domain to the Ndc80:Nuf2 coiled coil between the loop and tetramerization domains. See Material and Methods for details of how the model was obtained.

## DISCUSSION

The two structures reported here are both examples of using AF2 predictions to generate hypotheses for constructs likely to yield informative structures. For example, to design the particular loop construct that yielded the structure shown in Fig. 2D, ambiguities of coiled-coil heptad matching would have required a series of alternative pairings of the two chains on both sides of the loop itself, and the pairing C-terminal to the Ndc80 switchback is in fact half a heptad shifted from the best guess we had made previously, extrapolating from the structure of the four-way junction. AF2 predictions of the intervening coiled-coil region suggest how shifts in register (from the simple heptad-matched prediction) restore the experimentally determined pairing at the connection with Spc24:Spc25 (Valverde et al., 2016). The Dam1 segment docking would have been even harder to guess, as the successful design involved noticing a likely covalent linkage with Nuf2 (to ensure that the Dam1 fragment would be present in a crystal) and a deletion between the two ordered segments of Dam1. The structure also illustrates a modest limitation of the prediction, in the details of contacts between the short Dam1 α helix and the Ndc80:Nuf2 head (Fig. S4C).

The consistency of both the structure predictions and the final structures with genetic and biochemical data in the literature, as well as with each other, adds confidence to those predictions for which we do not yet have direct structural confirmation. For example, as illustrated in Fig. 2, the longest segment that could be deleted without lethality, in published experiments on the role of the loop in the transition to end-on microtubule attachment (Maure et al., 2011), coincides precisely with the region in Ndc80 of budding yeasts (*S. cerevisiae* and *K. lactis*, in Fig. 2) that appears in alignments as an “insertion” into Ndc80 loops of metazoans and even of fission yeast (see Data S1, Ndc80 sequence for *S. cerevisiae* starting at residue 490). Similarly, the AF2 predictions place Dam1 residues phosphorylated by Ipl1 at interface positions consistent with the physiological consequences of their modification.

We have integrated the information from the structures reported here with previously described analyses (Jenni et al., 2017; Jenni and Harrison, 2018) to generate the composite picture in Fig. 5. In addition to the switchback structure of the loop and the structurally defined interaction of the Dam1 segment with the Ndc80:Nuf2 globular domain, we have included additional contacts, based on inferences from biochemical and crosslinking mass-spectrometry data in the literature (Kim et al., 2017), conservation of segments in the flexible extensions of DASH/Dam1c components (Data S5 and S6), and sites at which phosphorylation affects microtubule attachment or detachment.

DASH/Dam1c binding at end-on attached kinetochores relies on oligomerization into rings, interactions with the microtubule lattice, and direct contacts with several Ndc80c that by themselves bind microtubules. Removal (by proteolysis with elastase) of the C-terminal “tails” of Dam1, Duo1 and Ask1 disables microtubule association of DASH/Dam1c heterodecamers *in vitro* as well as its polymerization into microtubule-encircling rings (Miranda et al., 2007). Additional experiments indicate that the tails of Dam1 and Duo1, but not of Ask1, are the principal microtubule contacts (Legal et al., 2016; Westermann et al., 2005). The negatively charged, flexible, C-terminal extensions of tubulin subunits are not required for DASH/Dam1c ring formation (Miranda et al., 2007). We therefore do not expect the microtubule contacts to be merely electrostatic compensation by positively charged residues but instead specific site-binding on the microtubule surface.

For Dam1, two short stretches of residues conserved among budding yeast are plausible candidates for the presumed microtubule contacts (Data S5). One is at the C-terminus of the Dam1 polypeptide chain (approximate residue range from 335 to 343); the other is in the middle of the extended segment between the structured, α-helical bundle at the core of the heterodecamer and the segment that binds the Ndc80:Nuf2 globular head (approximate residue range from 210 to 226). The latter includes conserved Mps1 phosphorylation sites serines 218 and 221, which appear to promote end-on attachment (Shimogawa et al., 2006). When the Ndc80:Nuf2 head is bound to a kinetochore microtubule, with a docked Dam1 segment, both candidate contacts will be close to the microtubule surface (Fig. 5). Duo1 and Dam1 emerge from the helical bundle of a DASH/Dam1c heterodecamer as a predicted, flexibly attached, approximately 30-residue, α-helical coiled coil (Dam1 residues 121–156, Duo1 residues 152– 179, both segments moderately well conserved among budding yeast, Data S5 and S6), allowing Duo1 residues 180 to 247 to project toward the microtubule. Conserved residues near the Duo1 C terminus (Data S6) are therefore also candidate microtubule contacts.

Interactions of DASH/Dam1c with Ndc80, defined by mass spectrometry cross-linking (Kim et al., 2017), include not only the segments of Dam1 included in the crystal structure in Fig. 3 (interaction A) but also segments in the C-terminal extensions of Ask1 (interaction B) and of the Spc19:Spc34 heterodimer that forms the protrusion domain of the DASH/Dam1c (interaction C). The crosslinked Ask1 segment centers on Ser200; phosphorylation of this residue by Ipl1 promotes dissociation (Flores et al., 2022). Cdc28/Cdk1 phosphorylation of Ask1 at serine 250, just at the edge of the crosslinked region, promotes DASH/Dam1c assembly on microtubules (Gutierrez et al., 2020; Higuchi and Uhlmann, 2005; Li and Elledge, 2003). C-terminal parts of Spc19:Spc34 together form the prominent protrusion that projects from the helical bundle at the core of the DASH/Dam1c heterodecamer. As suggested in Fig. 5, the protrusion is in a position to associate closely with the Ndc80:Nuf2 shaft between the loop and the tetramerization four-way junction. Cross links between residues in Spc34 and Ndc80 detected by mass spectrometry span the entire extent of the Spc34:Spc19 protrusion, centered on the Ipl1 phosphorylation site at Thr199 (Kim et al., 2017).

The Ndc80 loop participates in contacts essential for the transition from lateral to end-on kinetochore-microtubule attachment, in both yeast and human cells (Maure et al., 2011; Shrestha and Draviam, 2013). The six alanine substitutions in budding yeast Ndc80 that disrupt this transition (Maure et al., 2011) precisely span the disordered segment in the loop (Fig. 2). Because that segment is not essential for the structural integrity of Ndc80c, we infer that it probably binds some other kinetochore component or regulatory element. Although DASH/Dam1c is also essential for the end-on transition, the loop is not among the three contacts with Ndc80 defined by the published crosslinking experiments, discussed above, that motivated parts of the model in Fig. 5. The N-terminal extensions of Dam1 and Duo1 (60 and 57 residues, respectively), including an Ipl1 phosphorylation site at Dam1 serine 20 (Cheeseman et al., 2002), project together from the helical bundle of the heterodecamer relatively close to the loop (see asterisk in Fig. 5) and hence might be loop contacts.

In fission yeast, the properties of temperature-sensitive mutants in *S. pombe* Ndc80 suggest that the loop has a role in recruiting Dis1/TOG and Alp7/TACC-Alp14/TOG, orthologs of Stu2 (Hsu and Toda, 2011; Tang et al., 2013). In human cells, their ortholog, chTOG, also interacts with Hec1 (human Ndc80), at a yet-to-be-identified site, and like Stu2, has an Aurora-B independent role in microtubule-attachment error correction (Herman et al., 2020). Although the C-terminal segment of Stu2 binds the Ndc80c tetramerization domain, not the loop (Zahm et al., 2021), we cannot rule out additional functional contacts, either direct or indirect (Miller et al., 2019). Recent work presents evidence that the loop directs local clustering of human Ndc80c *in vitro* and shows that in cells, loop deletion or targeted mutation extends the spindle assembly checkpoint and delays anaphase (Polley et al., 2022). Human cells lack DASH/Dam1c, however, and recruit the non-homologous Ska complex instead (van Hooff et al., 2017), and their centromeres bind bundles of at least 10 microtubules (Kiewisz et al., 2022; O’Toole et al., 2020), rather than just one, probably with higher Ndc80c density than in yeast cells (Suzuki et al., 2015). The correspondence of predicted loop structures from metazoans and fission yeast, whose centromeres bind several microtubules, suggests that the loop might have a role in coordinating interaction with multiple microtubules. Thus, despite similar functional outcomes, not all the molecular interactions in loop-directed, end-on binding need be homologous in metazoans and budding yeast.

## MATERIALS AND METHODS

### AF2 predictions

For the AlphaFold predictions, we ran AF2, implemented either by SBgrid (version 2.1.1) or the O2 cluster at Harvard Medical School (version 2.2.2).

### Ndc80c loop x-ray structure determination

To express the loop region of human or yeast Ndc80c, as a recombinant heterodimer, we cloned codon-optimized synthetic genes (IDT) coding for Ndc80 (residues 370–509 for human Ndc80, Data S1) with a TEV protease-cleavable 6-His tag and Nuf2 (residues 252–347 for human Ndc80, Data S2) into the respective multiple cloning sites of pETDuet-1. To facilitate formation of the Ndc80/Nuf2 heterodimer, we introduced four cysteine residues, two on the respective N and C termini of Ndc80, and another two on those of Nuf2. Proteins were co-expressed in Shuffle T7 *E. coli* (New England Biolabs). Low yield of the expressed yeast protein led us to focus on the human complex. We grew a total of 6 liters of culture in 12 2 L baffled flasks at 37 °C with shaking at 225 rpm to an O.D. of 0.8. Cells were then induced with 200 µM IPTG and incubated overnight at 18 °C with shaking at 225 rpm. Cells were harvested by centrifugation and resuspended in 150 mL of a buffer containing 50 mM TRIS pH 8.0, 250 mM NaCl, 10 mM imidazole pH 8.0, 5 mM β-mercaptoethanol, 1 mM PMSF, 1 µg/mL pepstatin, 1 µg/mL aprotinin, 1 µg/mL leupeptin, 83 µg/mL lysozyme, and 30 µg/mL DNase I. Cells were lysed by sonication, and the lysate was clarified by centrifugation and applied to Ni-NTA agarose equilibrated with buffer containing 20 mM TRIS pH 8.0, 100 mM NaCl, 10 mM imidazole pH 8.0, and 5 mM β-mercaptoethanol (equilibration buffer). Bound material was washed with equilibration buffer containing 20 mM imidazole pH 8.0 and 500 mM NaCl and then with equilibration buffer. Proteins were eluted with equilibration buffer containing 400 mM imidazole pH 8.0 and 50 mM NaCl. Eluates were treated overnight with TEV protease to remove the 6-His tag from the N terminus of Ndc80 and subjected to anion exchange chromatography using a HiTrap Q HP column (Cytiva), followed by size-exclusion chromatography using a HiLoad Superdex 200 26/60 column (Cytiva) equilibrated with 10 mM HEPES pH 7.5, 100 mM NaCl, and 50 mM β-mercaptoethanol.

The human Ndc80c loop complex was concentrated to 25 mg/mL in a in an Amicon centrifugal concentrator in a buffer containing 10 mM HEPES pH 7.5, 100 mM NaCl, 2 mM TCEP, and 50 mM β-mercaptoethanol. The protein complex was crystallized by hanging drop vapor diffusion using 1 µL protein solution and 1 µL of a well solution containing 20% (w/v) polyethylene glycol 4000, 0.1 M TRIS pH 8.6, and 0.2 M magnesium chloride. Crystals were cryo-protected by a 3–5 min soak in well solution supplemented with 25% (v/v) glycerol and flash-frozen in liquid nitrogen.

The complex crystallized in in space group C_2_ (a = 49.9 Å, b = 72.3 Å, c = 80.1 Å). Data to a minimum Bragg spacing of 1.9 Å were recorded on the NE-CAT beamline 24-ID-C at the Advanced Photon Source (APS), and indexed, integrated, scaled, and merged using the dials-aimless pipeline in xia2 (Gildea et al., 2022) (Table S1). The structure was determined by molecular replacement using Phaser (McCoy et al., 2007) as implemented in CCP4 Cloud (Krissinel et al., 2022). Search models consisted of four fragments of the AF2-generated structure. Model building was carried out in Coot (Emsley et al., 2010), and refinement, in Phenix (Adams et al., 2010). We used MolProbity (Chen et al., 2010) for structure validation. The final model includes the following residues (Table S2 and Data S1–S2): Ndc80^370–509^, Nuf2^252–347^, plus one cysteine residue at both ends of each chain and one serine (left after proteolysis by TEV) at the N terminus of Ndc80. Model statistics are shown in Table S1.

### Ndc80:Dam1-Nuf2 x-ray structure determination

We designed an Ndc80/Dam1-Nuf2 fusion construct, with sequences from *S. cerevisiae*, based on the AlphaFold prediction, as described in the main text. To express this construct as recombinant protein, we modified a previously constructed pETDuet vector coding for truncated (“dwarf”) versions of Ndc80 and Nuf2 (Valverde et al., 2016; Zahm et al., 2021), by fusing segments from the C-terminal extension of Dam1 (residues 252–270 and 290–305) to the N terminus of Nuf2 (Data S3–S5). To express a Dam1-fused heterodimer, we co-transformed Rosetta2(DE3)pLysS with the modified construct and a pRSFDuet plasmid containing the genes for truncated dwarf versions of Spc24 and Spc25 (Valverde et al., 2016; Zahm et al., 2021). We grew a total of 6 liters of culture in 12 2 L baffled flasks at 37 °C with shaking at 225 rpm to an O.D. of 0.8. Cells were then induced with 200 µM IPTG, followed by overnight incubation at 18 °C with shaking at 225 rpm. The Dam1-fused, dwarf Ndc80c was purified as described above for the recombinant loop region of human Ndc80c (see above), except that the buffer of the final gel filtration step contained 2 mM tris(2-carboxyethyl)phosphine (TCEP) instead of β-mercaptoethanol.

The dwarf-Ndc80c with the Dam1 segment fused to the N terminus of Nuf2 was concentrated to 15 mg/mL in an Amicon centrifugal concentrator in a buffer containing 10 mM HEPES pH 7.5, 100 mM NaCl, and 2 mM TCEP. The protein complex was crystallized by hanging drop vapor diffusion using 1 µL protein solution and 1 µL of a well solution containing 12% (v/v) 2-methyl-2,4-pentanediol (MPD) and 0.1 M ADA buffer pH 6.8. Crystals were cryoprotected by soaking for 3– 5 min in well solution containing 25% (v/v) MPD followed by flash-freezing in liquid nitrogen.

Diffraction data from multiple crystals were recorded on the NE-CAT beamline 24-ID-E at the Advanced Photon Source (APS), each with a total range of 360°. Data were indexed, integrated, scaled, and merged using the DIALS-aimless pipeline (Winn et al., 2011; Winter et al., 2022) as implemented in xia2 (Gildea et al., 2022). To achieve higher redundancy and thus get better estimates of the intensities of weak reflections, 9 isomorphous datasets were scaled together and merged. Indexing of the diffraction pattern showed a primitive hexagonal Bravais lattice for the crystallized complex. Scaling and molecular replacement (see below) showed that the space group was P3_2_12 (a = b = 130.0 Å, c = 216.3 Å) (Table S1).

The structure was determined by molecular replacement using Phaser (McCoy et al., 2007) as implemented in Phenix (Adams et al., 2010). The search model consisted of the Ndc80 and Nuf2 heads with a stretch of coiled coil. We could not place any search models that contained Spc24:Spc25, nor did we observe any density for Spc24:Spc25 in difference maps. From the placement of Ndc80:Nuf2 it was evident that the crystal packing did not allow the presence of Spc24:Spc25 as seen in the dwarf-Ndc80c structure (Valverde et al., 2016; Zahm et al., 2021). Thus, during crystallization, Spc24:Spc25 had dissociated from Ndc80:Dam1-Nuf2, and there were two copies of a Ndc80:Dam1-Nuf2 complex in the asymmetric unit.

To obtain structure factors for model building, phase improvement and structure refinement, we submitted unmerged data after integration with xia2 (Gildea et al., 2022) to the STARANISO server (Tickle et al., 2018). After excluding frames with high merging R values, we used the “unmerged data” protocol for Bayesian estimation of structure amplitudes. We kept the default value of 1.20 for the local mean I/σ(I) as diffraction cut-off criterion, which resulted in diffraction limits at the principal axes of the ellipsoid fitted to the diffraction cut-off surface of 3.71 Å (principal axis 0.894 **a\*** - 0.447 **b\***), 3.71 Å (principal axis **b\***), and 3.14 Å (principal axis **c\***). Final merging statistics from the STARANISO processing are given in Table S2. We used uniquify from CCP4 (Winn et al., 2011) to select 5% of the reflections for the calculation of free R factors and B-sharpened the observed amplitudes with phenix.remove_aniso (b_iso = 80.0).

To obtain the best possible, unbiased electron density for the Dam1 segment, we modeled and refined Ndc80:Nuf2 to convergence before modeling the Dam1 segment. For this modeling, we manually adjusted the Ndc80 and Nuf2 models in O (Jones et al., 1991) and Coot (Emsley et al., 2010), followed by refinement with phenix.refine (Afonine et al., 2012), and density modification with RESOLVE (Terwilliger, 2000). We ran RESOLVE with 2-fold non-crystallographic symmetry (NCS) averaging, restricted to regions occupied by Ndc80 and Nuf2. We defined 5 NCS groups (Ndc80: residues 114–237, 238–626, 627–684; Nuf2: residues 13–136, 137–451) and calculated transformation matrices and masks based on the corresponding structures. Because the density corresponding to the Dam1 segment differed substantially between the two NCS copies (see below), this region was excluded from NCS averaging. We also outlined the Dam1 segment density by placing dummy atoms, which we then used to exclude this region from the solvent mask in RESOLVE. We input a solvent content fraction of 0.72. We also calculated feature-enhanced maps with phenix.fem (Afonine et al., 2015). An electron density map from the last iteration of model building, refinement and phase improvement before modeling the Dam1 segment and a final map are shown in Figure S3.

We observed that density for the Dam1 segment fused to the second copy of the Ndc80:Nuf2 head in the asymmetric unit was much weaker than the first, and its appearance was of insufficient clarity to build a molecular model. Superposition of our Ndc80:Dam-Nuf2 structure from NCS position 1 onto Ndc80:Nuf2 of NCS position 2 showed that a Dam1 segment bound as observed at position 1 would clash at position 2 with its crystallographic symmetry mate. Thus, the crystal packing prevented uniform ordering of the Dam1 segment at position 2.

We refined final structure with phenix.refine (Afonine et al., 2012) against B-sharpened structure amplitudes obtained from STARANISO. The final model includes the following residues (Table S2 and Data S3–S5): Ndc80^114–318, 621–684^ (chain A), Nuf2^2–153, 407–451^ (chain B), Dam1^254–270, 290–305^ (chain B), Ndc80^115–318, 621–684^ (chain C), Nuf2^9–153, 407–451^ (chain D). In addition to standard stereochemical restraints, we used secondary structure, Ramachandran, and torsion angle NCS restraints. Target function weight optimization was turned on. For grouped anisotropic B factor refinement, TLS groups were obtained from the TLS Motion Determination (TLSMD) web server (Painter and Merritt, 2006) (Table S2). We used MolProbity (Chen et al., 2010) for structure validation. The final model statistics are shown in Table S1.

### Model of the yeast kinetochore-microtubule interface

We modeled a 13-protofilament MT based on the cryoEM structure of a yeast tubulin dimer polymerized with GTP *in vitro* (PDB-ID 5W3F) (Howes et al., 2017). As published, we used lattice parameters for the MT tube with a helical rise of 83.3 Å and a rotation of 0.43° between dimers within a protofilament, and a helical rise of 9.65 Å and a rotation of −27.6° between protofilaments. The majority of kinetochore MTs in tomograms recorded from budding yeast were observed in in “ram’s horn” geometry (McIntosh et al., 2013), and we modeled the flaring of protofilaments at the MT plus end by matching the average curvature of the tomograms with the curvature observed in various structures of bended tubulin protofilaments (PDB-IDs 3J6H, 3RYH, 4HNA, 4FFB, 6MZG).

To model a MT-bound yeast Ndc80c, we first docked the AF2 prediction of Ndc80:Nuf2 up to a few residues after the hinge (Ndc80^115–444^, Nuf2^1–277^) onto the MT. For this, we used the structure of the human Ndc80:Nuf2 head domain bound to the MT lattice (PDB-ID 3iZ0) (Alushin et al., 2010) for superposition of tubulin and the Ndc80 head domain (*H. sapiens* residues 110–202). Next, we placed a composite model of two AlphaFold 2 predictions, comprising sequence from Ndc80:Nuf2 just before the hinge all the way to the Spc24:Spc25 head domains (Ndc80^413–691^, Nuf2^252–451^, Spc25^1–89^, Spc25^1–83^; and Ndc80^619–691^, Nuf2^404–451^, Spc24^1–213^, Spc25^1–221^) with a hinge angle such that the coiled coils of Ndc80:Nuf2 (after the hinge) and Spc24:Spc25 were approximately parallel to the microtubule axis. Finally, we re-modeled the hinge residues in plausible conformation using RosettaRemodel (Huang et al., 2011).

For the DASH/Dam1c, we manually placed one heterodecamer of the *S. cerevisiae* full-length AlphaFold 2 prediction, including only the well-structured regions of the core complex with high confidence scores (Ask1^2–69^, Dad1^14–73^, Dad2^2–85, 116–133^, Dad3^6–94^, Dad4^2–72^, Dam1^54–162^, Duo1^61–180^, Hsk3^2–69^, Spc19^2–106^, Spc34^2–118,157–264^), into the cryoEM map of a DASH/Dam1c ring assembled around a MT (Ramey et al., 2011) (note that the deposited map, EMD-5254, has the wrong hand and needs to be inverted), followed by rigid-body fitting with phenix.real_space_refine (Afonine et al., 2018). The full-length complex was then placed on the fitted core complex and 17-fold rotationally expanded around the MT axis. The whole DASH/Dam1c ring was then rotated around and translated along the MT axis such that interaction C (Flores et al., 2022; Kim et al., 2017) between the protrusion domain of one DASH/Dam1c heterodecamer and the MT-bound Ndc80c could be established. This juxtaposed Thr199 of Spc34, the residue that is phosphorylated by Ipl1 and regulates interaction C (Flores et al., 2022), and residue 583 of Ndc80, the position of a five amino acid mutation (insertion) that abrogates interaction C (Kim et al., 2017). It also allows for establishment of the interaction between Spc19 residues 128–165 and Nuf2 residues 399–429 (Wang et al., 2012). The C termini of Spc19 and Sp34 form a coiled coil at the tip of the DASH/Dam1c protrusion domain. It was not observed in the cryoEM reconstruction of the *C. thermophilum* DASH/Dam1c complex (because of its flexible attachment), but it was inferred from sequence analysis (Jenni and Harrison, 2018) and AlphaFold 2 also predicts it now. We used HADDOCK 2.4 (van Zundert et al., 2016) to dock the C-terminal Spc19:Spc34 coiled coil onto Ndc80:Nuf2 and re-modeled the residues that connect it to the protrusion domain with RosettaRemodel (Huang et al., 2011). The model of the DASH/Dam1c ring shown in Fig. 5 contains the following residues for each subunit: Ask1^1–70^, Dad1^15–77^, Dad2^1–73, 119–133^, Dad3^6–35, 49–94^, Dad4^3–70^, Dam1^53–156^, Duo1^58–179^, Hsk3^3–66^, Spc19^1–165^, Spc34^1–295^. We added the C-terminal Dam1 segment (residues 254–270, 290–305) from the crystal structure determined here by superposition of the Ndc80 head domains.

## ACKNOWLEDGMENTS

We thank the staff of the NE-CAT beamlines, in particular Jon Scheurman, for help in data collection. NE-CAT is supported by NIH grant P30 GM124165, using resources of the Advanced Photon Source, operated by Argonne National Laboratory under Contract DE-AC02-06CH11357. Portions of this research were conducted on the O2 High Performance Compute Cluster, operated by the Research Computing Group at Harvard Medical School. SCH is an Investigator in the Howard Hughes Medical Institute.

---

**Fig. S1.**
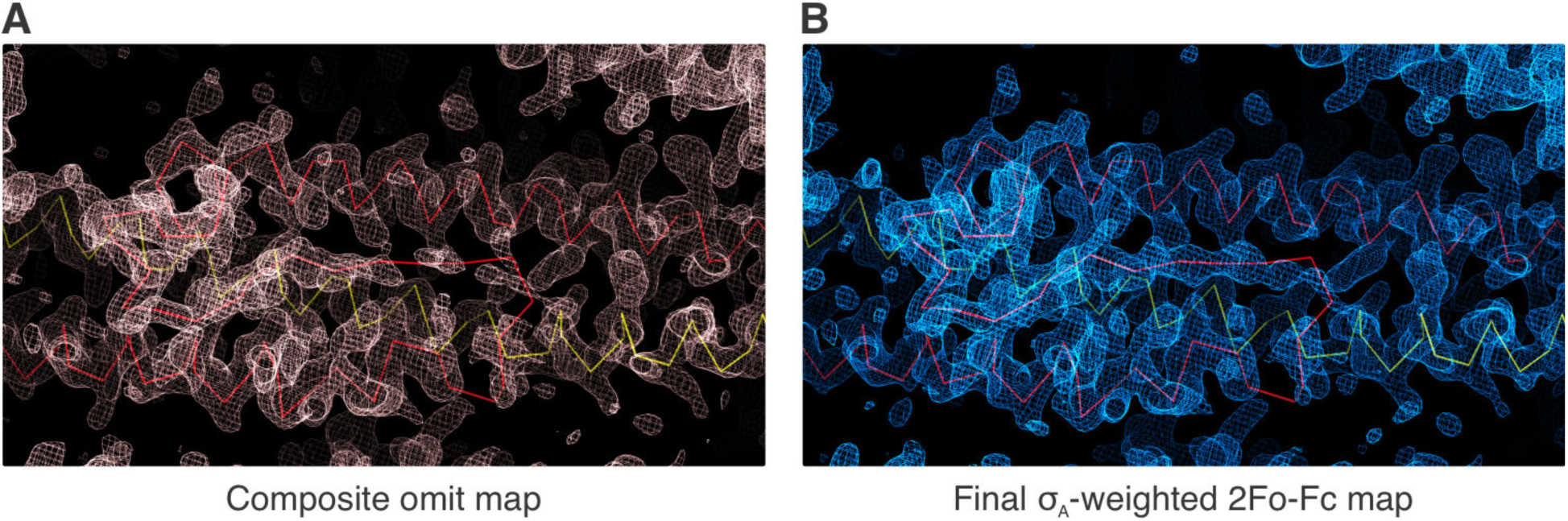
Ndc80 loop (human) electron density maps. A Cα-ribbon trace of the final model is shown with Ndc80 (red), Nuf2 (yellow). (A) Composite omit map shown as pink mesh. (B) Final α_A_-weighted 2mF_o_-DF_c_ difference map shown as blue mesh.

**Fig. S2.**
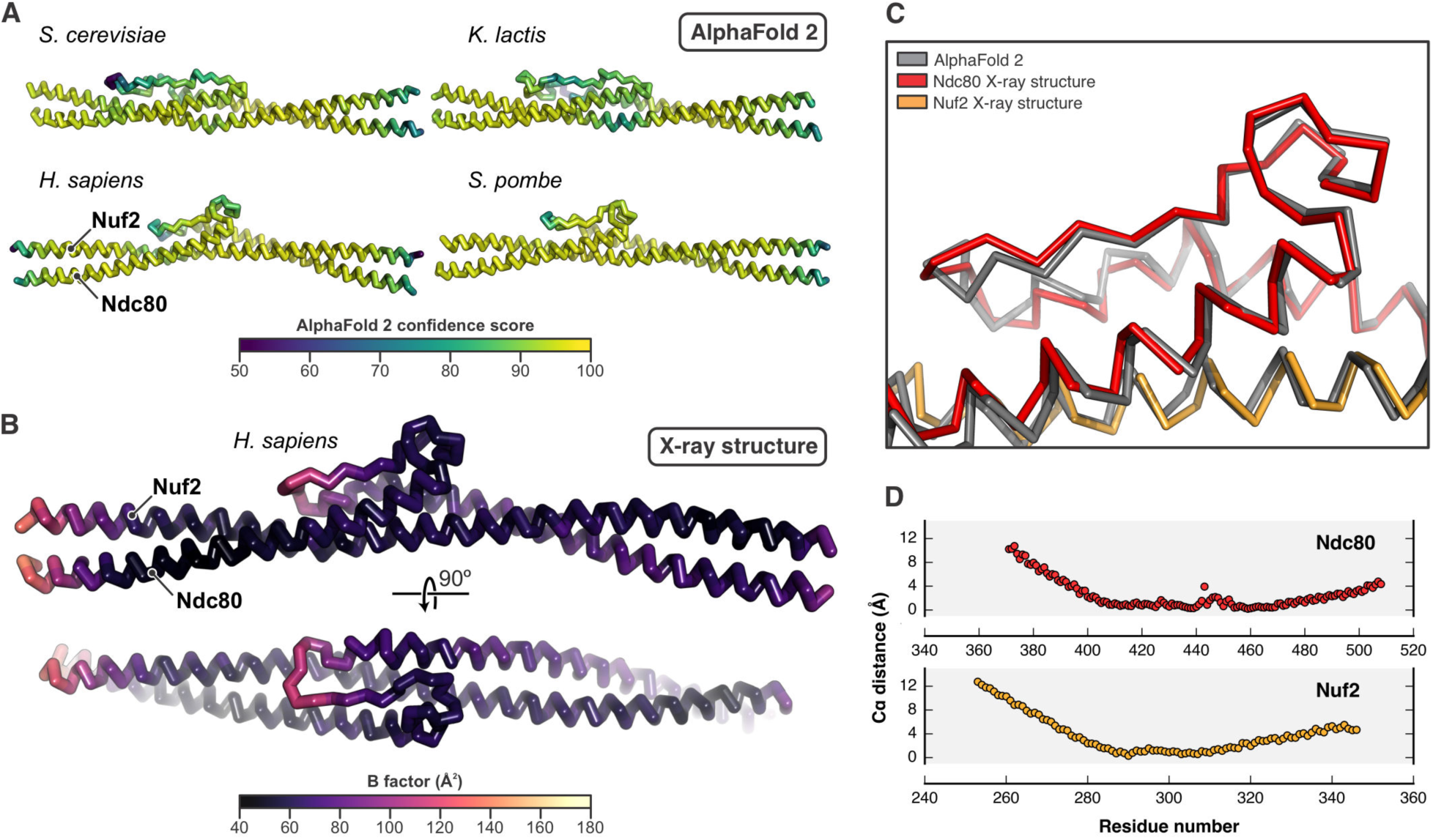
Comparison of the Ndc80 loop AF2 predictions and the crystal structure. (A) The AF2 predictions from different species as shown in Fig. 2C are in Cα ribbon representations and colored according to the AF2 per-residue confidence scores. (B) The crystal structure of the human Ndc80 loop as shown in Fig. 2D is in Cα ribbon representation and colored according to its B factors. (C) Close-up view of the loop region of the superposition of the human AF2 prediction (gray) onto the crystal structure (colored) (residues for superposition: Ndc80, 402–478; Nuf2, residues 282–317). (D) Per-residue Cα distances between the human AF2 prediction and the crystal structure after superposition (residues for superposition: Ndc80, 402–478; Nuf2, residues 282–317).

**Fig. S3.**
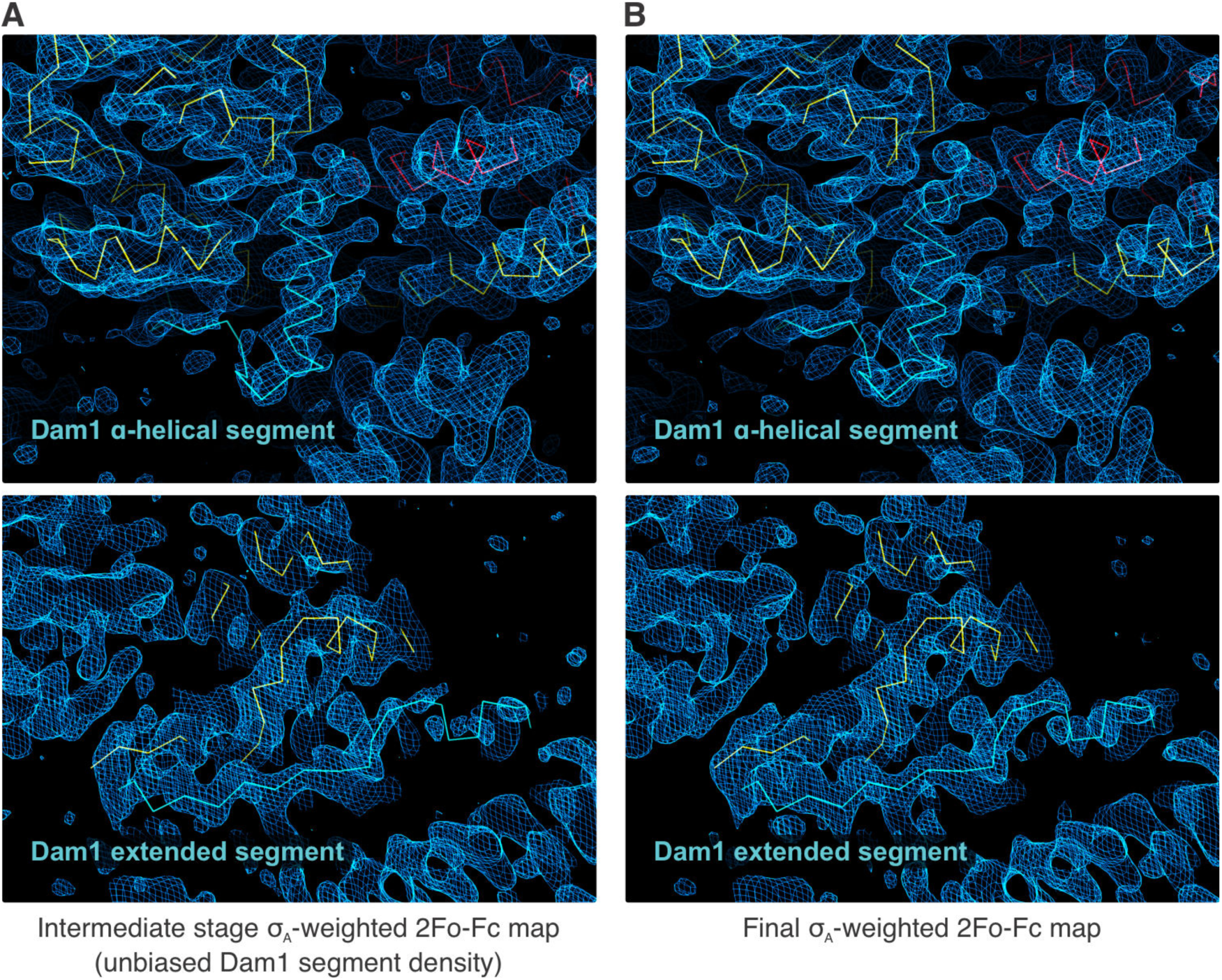
Ndc80:Dam1-Nuf2 (yeast) electron density maps. A Cα-ribbon trace of the final model is shown with Ndc80 (red), Nuf2 (yellow), and Dam1 (cyan). The electron density maps are shown as blue mesh. (A) Intermediate stage α_A_-weighted 2mF_o_-DF_c_ difference map. The density of the Dam1 segment is unbiased in this map, because it was not included in the model up to this stage. (B) Final α_A_-weighted 2mF_o_-DF_c_ difference map.

**Fig. S4.**
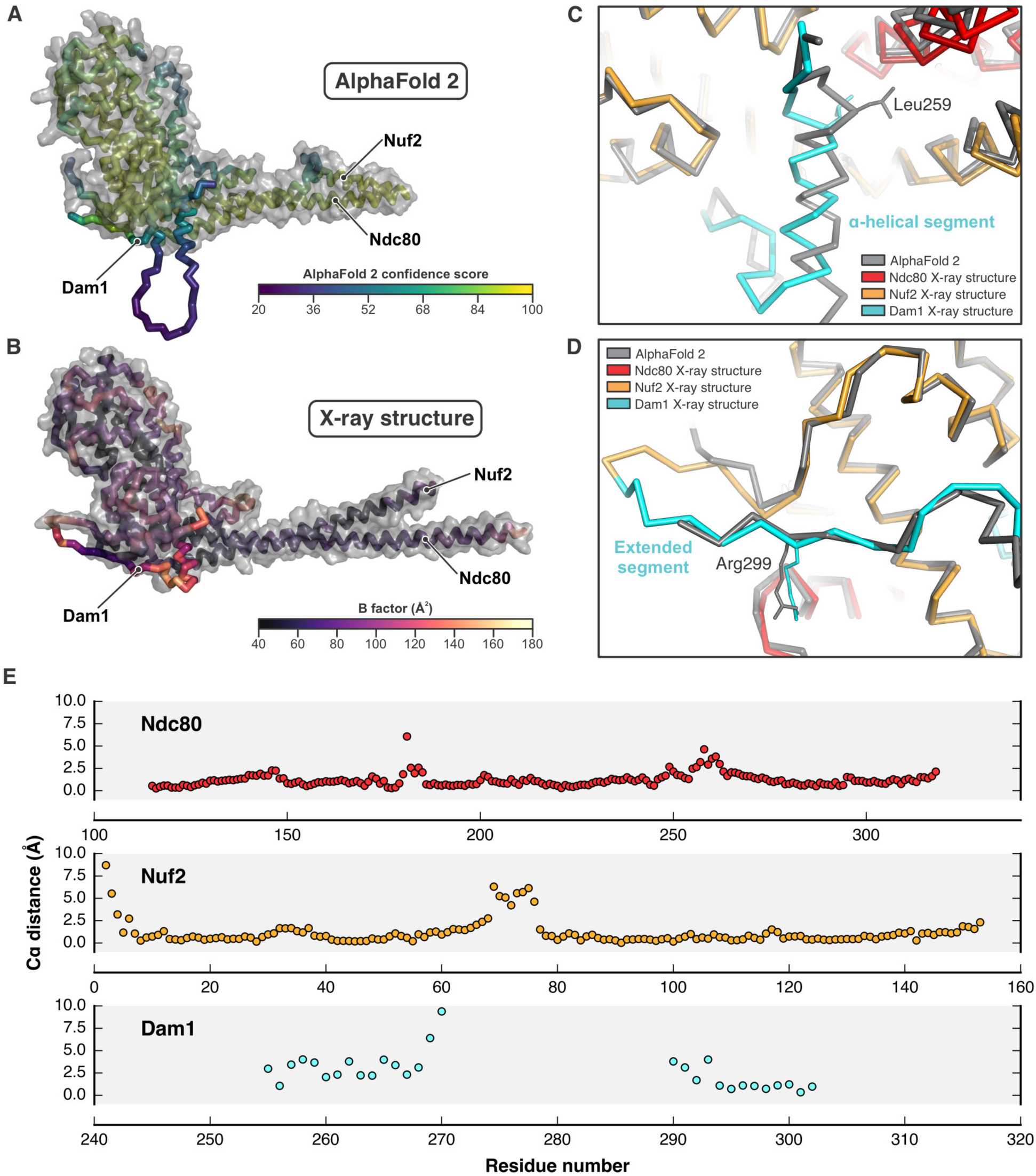
Comparison of Ndc80:Nuf2:Dam1 AF2 prediction and the crystal structure. (A) The AF2 prediction as shown in Fig. 3B is in Cα ribbon representation with transparent surface for Ndc80 and Nuf2, and colored according to the AF2 per-residue confidence scores. (B) The crystal structure of the yeast Ndc80:Nuf2:Dam1 complex as shown in Fig. 3C is in Cα ribbon representation with transparent surface for Ndc80 and Nuf2, and colored according to its B factors. (C) Superposition of the AF2 prediction (gray) onto the crystal structure (colored). A close-up view of the Dam1 α-helical segment is shown. (C) Superposition of the AF2 prediction (gray) onto the crystal structure (colored). A close-up view of the Dam1 extended segment is shown. (E) Per-residue Cα distances between the AF2 prediction and the crystal structure after superposition.

**Table S1.**
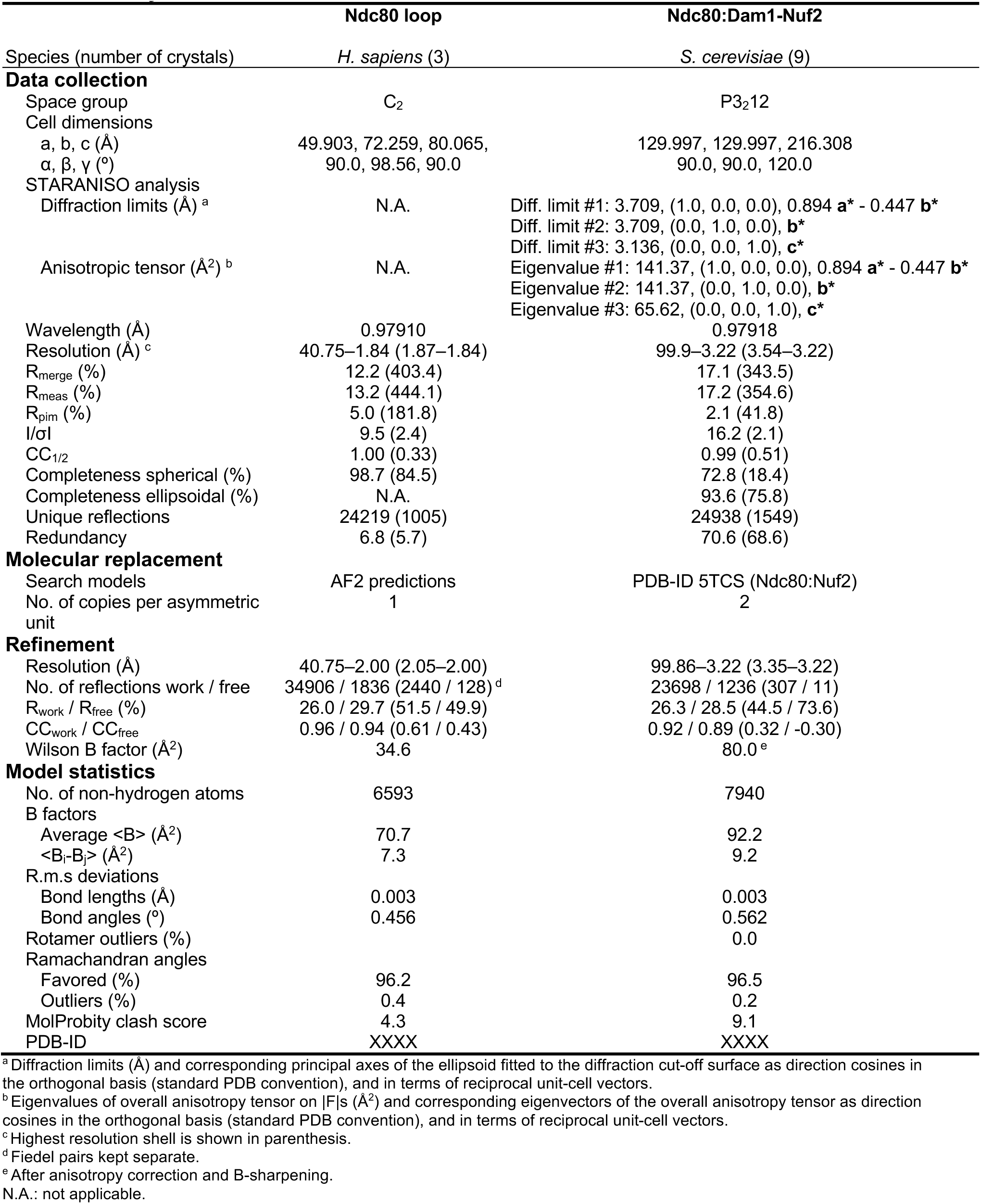
X-ray data collection and structure refinement statistics.

|  | Ndc80 loop | Ndc80:Dam1-Nuf2 |
| --- | --- | --- |
| Species (number of crystals) | *H. sapiens* (3) | *S. cerevisiae* (9) |
| **Data collection** |  |  |
| Space group | C<sub>2</sub> | P3<sub>2</sub>12 |
| Cell dimensions |  |  |
| a, b, c (Å) | 49.903, 72.259, 80.065, | 129.997, 129.997, 216.308 |
| α, β, γ (°) | 90.0, 98.56, 90.0 | 90.0, 90.0, 120.0 |
| STARANISO analysis |  |  |
| Diffraction limits (Å)<sup>a</sup> | N.A. | Diff. limit #1: 3.709, (1.0, 0.0, 0.0), 0.894 **a\*** - 0.447 **b\***<br>Diff. limit #2: 3.709, (0.0, 1.0, 0.0), **b\***<br>Diff. limit #3: 3.136, (0.0, 0.0, 1.0), **c\*** |
| Anisotropic tensor (Å<sup>2</sup>)<sup>b</sup> | N.A. | Eigenvalue #1: 141.37, (1.0, 0.0, 0.0), 0.894 **a\*** - 0.447 **b\***<br>Eigenvalue #2: 141.37, (0.0, 1.0, 0.0), **b\***<br>Eigenvalue #3: 65.62, (0.0, 0.0, 1.0), **c\*** |
| Wavelength (Å) | 0.97910 | 0.97918 |
| Resolution (Å)<sup>c</sup> | 40.75–1.84 (1.87–1.84) | 99.9–3.22 (3.54–3.22) |
| R<sub>merge</sub> (%) | 12.2 (403.4) | 17.1 (343.5) |
| R<sub>meas</sub> (%) | 13.2 (444.1) | 17.2 (354.6) |
| R<sub>pim</sub> (%) | 5.0 (181.8) | 2.1 (41.8) |
| I/σI | 9.5 (2.4) | 16.2 (2.1) |
| CC<sub>1/2</sub> | 1.00 (0.33) | 0.99 (0.51) |
| Completeness spherical (%) | 98.7 (84.5) | 72.8 (18.4) |
| Completeness ellipsoidal (%) | N.A. | 93.6 (75.8) |
| Unique reflections | 24219 (1005) | 24938 (1549) |
| Redundancy | 6.8 (5.7) | 70.6 (68.6) |
| **Molecular replacement** |  |  |
| Search models | AF2 predictions | PDB-ID 5TCS (Ndc80:Nuf2) |
| No. of copies per asymmetric unit | 1 | 2 |
| **Refinement** |  |  |
| Resolution (Å) | 40.75–2.00 (2.05–2.00) | 99.86–3.22 (3.35–3.22) |
| No. of reflections work / free | 34906 / 1836 (2440 / 128)<sup>d</sup> | 23698 / 1236 (307 / 11) |
| R<sub>work</sub> / R<sub>free</sub> (%) | 26.0 / 29.7 (51.5 / 49.9) | 26.3 / 28.5 (44.5 / 73.6) |
| CC<sub>work</sub> / CC<sub>free</sub> | 0.96 / 0.94 (0.61 / 0.43) | 0.92 / 0.89 (0.32 / -0.30) |
| Wilson B factor (Å<sup>2</sup>) | 34.6 | 80.0<sup>e</sup> |
| **Model statistics** |  |  |
| No. of non-hydrogen atoms | 6593 | 7940 |
| B factors |  |  |
| Average <B> (Å<sup>2</sup>) | 70.7 | 92.2 |
| <B<sub>i</sub>-B<sub>j</sub>> (Å<sup>2</sup>) | 7.3 | 9.2 |
| R.m.s deviations |  |  |
| Bond lengths (Å) | 0.003 | 0.003 |
| Bond angles (°) | 0.456 | 0.562 |
| Rotamer outliers (%) |  | 0.0 |
| Ramachandran angles |  |  |
| Favored (%) | 96.2 | 96.5 |
| Outliers (%) | 0.4 | 0.2 |
| MolProbity clash score | 4.3 | 9.1 |
| PDB-ID | XXXX | XXXX |

<sup>a</sup> Diffraction limits (Å) and corresponding principal axes of the ellipsoid fitted to the diffraction cut-off surface as direction cosines in the orthogonal basis (standard PDB convention), and in terms of reciprocal unit-cell vectors.
<sup>b</sup> Eigenvalues of overall anisotropy tensor on |F|s (Å<sup>2</sup>) and corresponding eigenvectors of the overall anisotropy tensor as direction cosines in the orthogonal basis (standard PDB convention), and in terms of reciprocal unit-cell vectors.
<sup>c</sup> Highest resolution shell is shown in parenthesis.
<sup>d</sup> Fiedel pairs kept separate.
<sup>e</sup> After anisotropy correction and B-sharpening.
N.A.: not applicable.

**Table S2.**
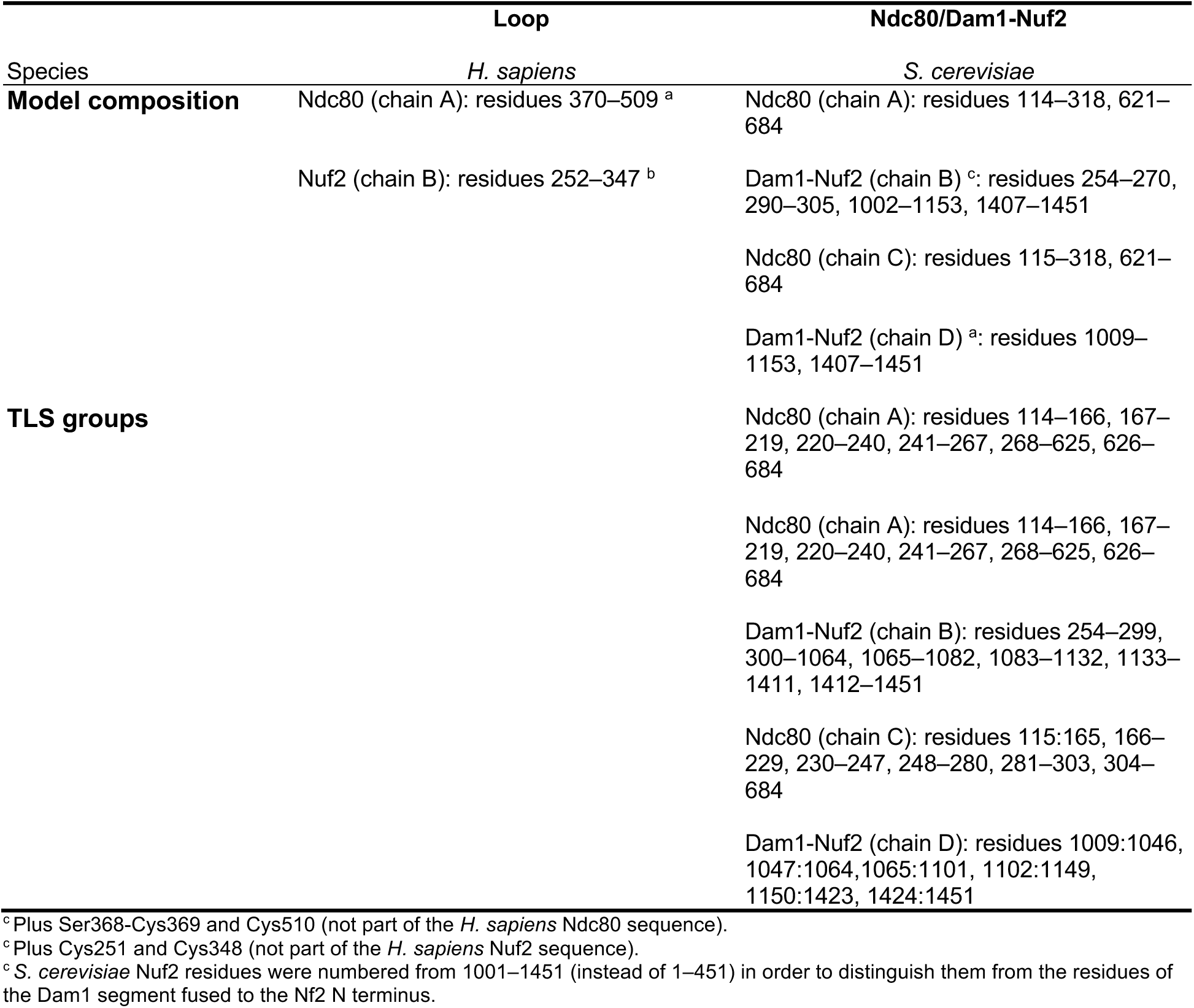
Model compositions and TLS groups.

## SUPPLEMENTARY DATA

**Data S1.**
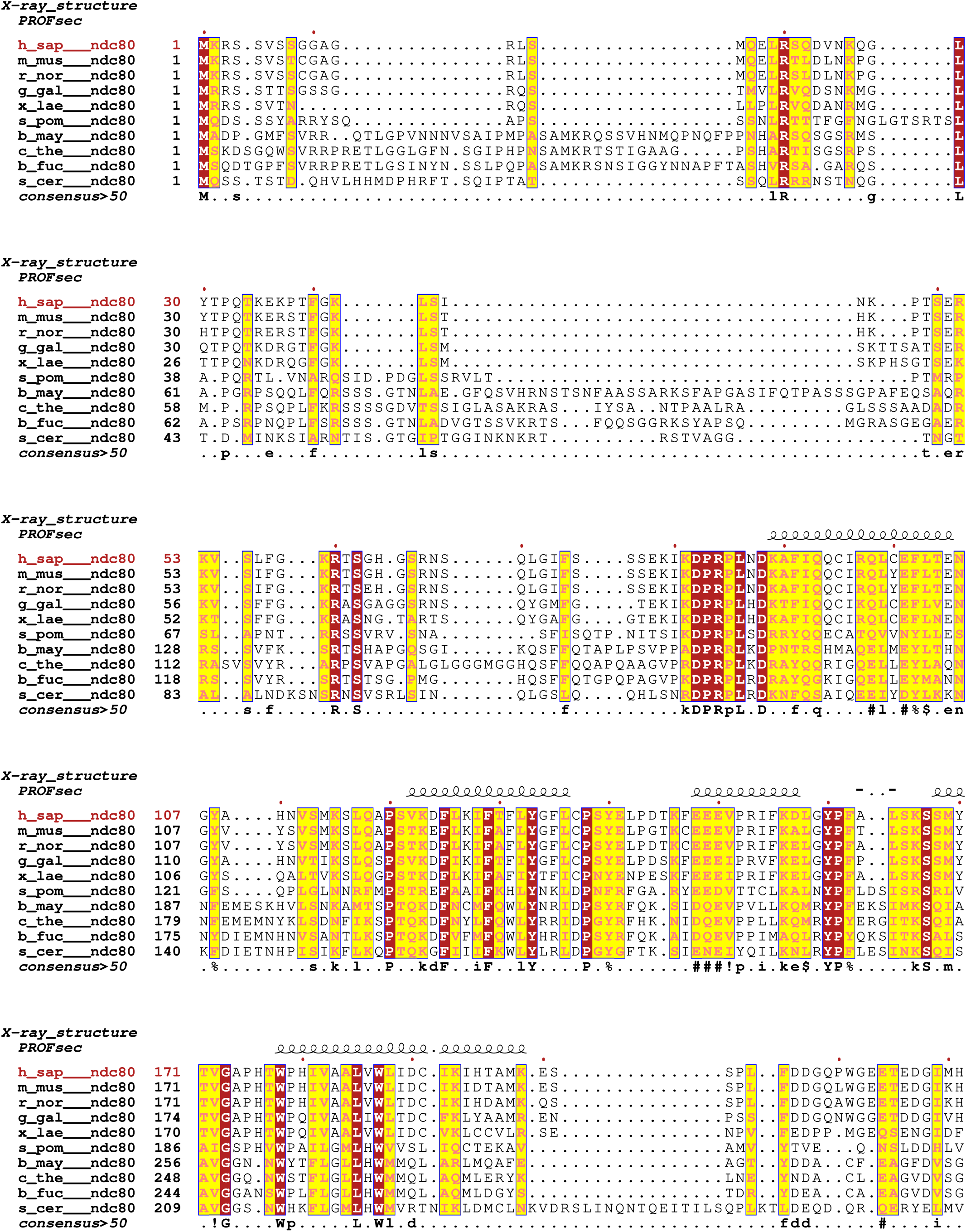
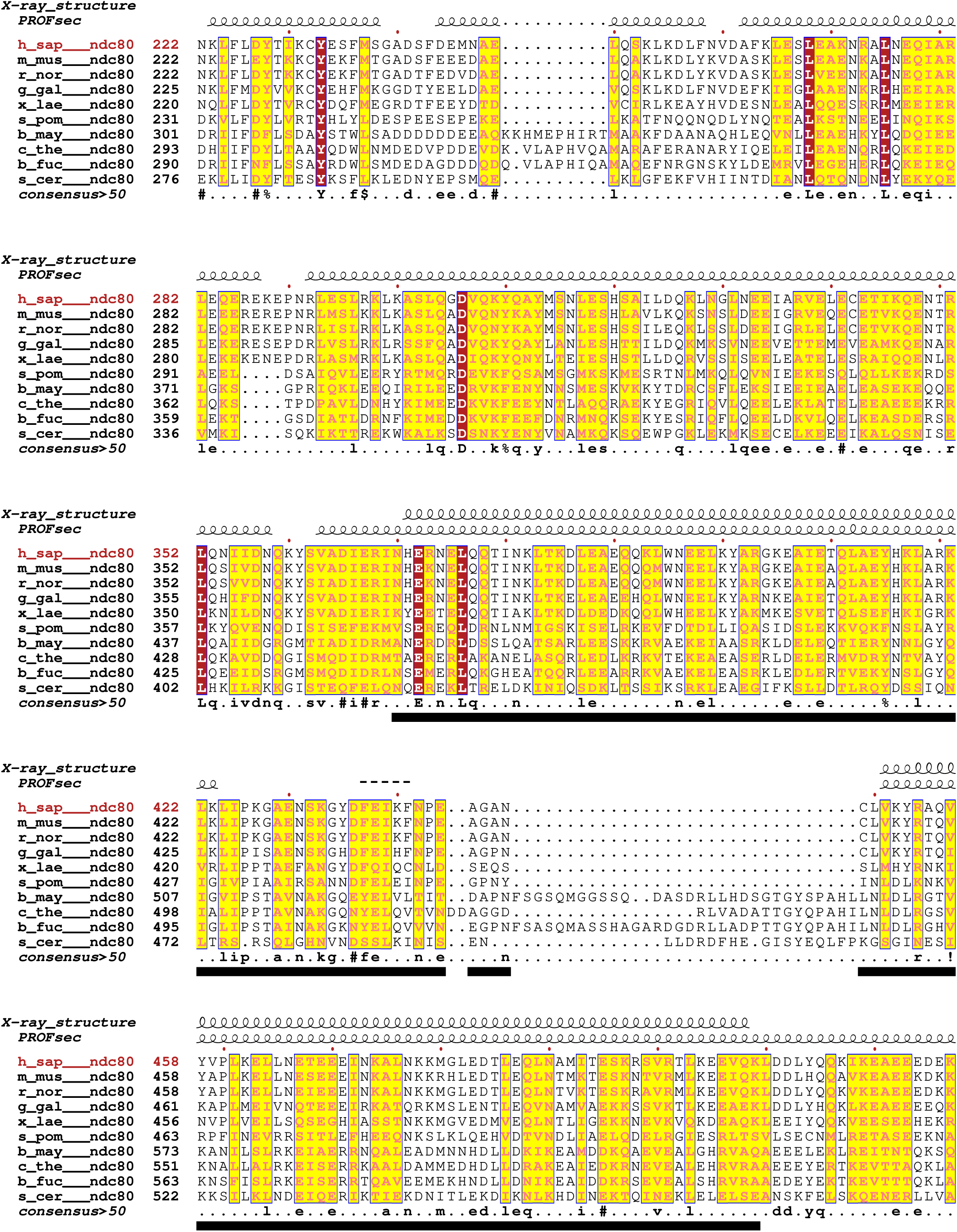
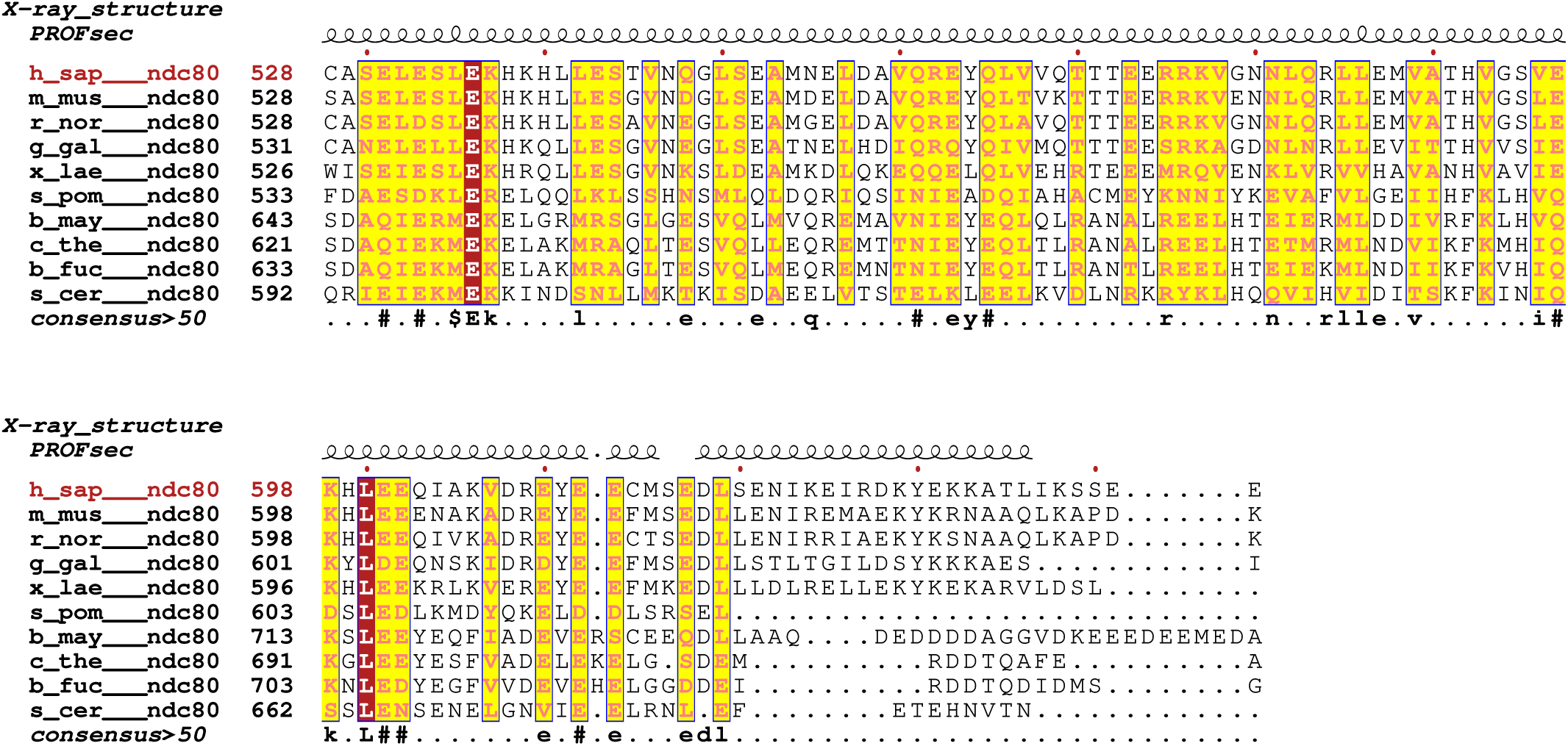
Ndc80 multiple-sequence alignment from metazoans and fungi. Sequences were retrieved from the UniProt (UniProt, 2021) database with the following accession identifiers: h_sap: *Homo sapiens*, O14777; m_mus: *Mus musculus*, Q9D0F1; r_nor: *Rattus norvegicus*, F7F189; g_gal: *Gallus gallus*, Q76I89; x_lae: *Xenopus laevis*, Q8AWF5; s_pom: *Schizosaccharomyces pombe*, Q10198; b_may: *Bipolaris maydis*, M2UME3; c_the: *Chaetomium thermophilum*, G0S8W8; b_fuc: *Botryotinia fuckeliana*, M7U0R4; s_cer: *Saccharomyces cerevisiae*, P40460. Sequences were aligned with T-Coffee (Notredame et al., 2000) and displayed with ESPript (Robert and Gouet, 2014). Above the sequences, secondary structure is shown as annotated by DSSP (Kabsch and Sander, 1983) from the Ndc80:Nuf2 loop crystal structure or as predicted with PROFsec (Rost, 2001), respectively. The black bars below the sequences are residues that were modeled in the Ndc80:Nuf2 loop crystal structure here.

**Data S2.**
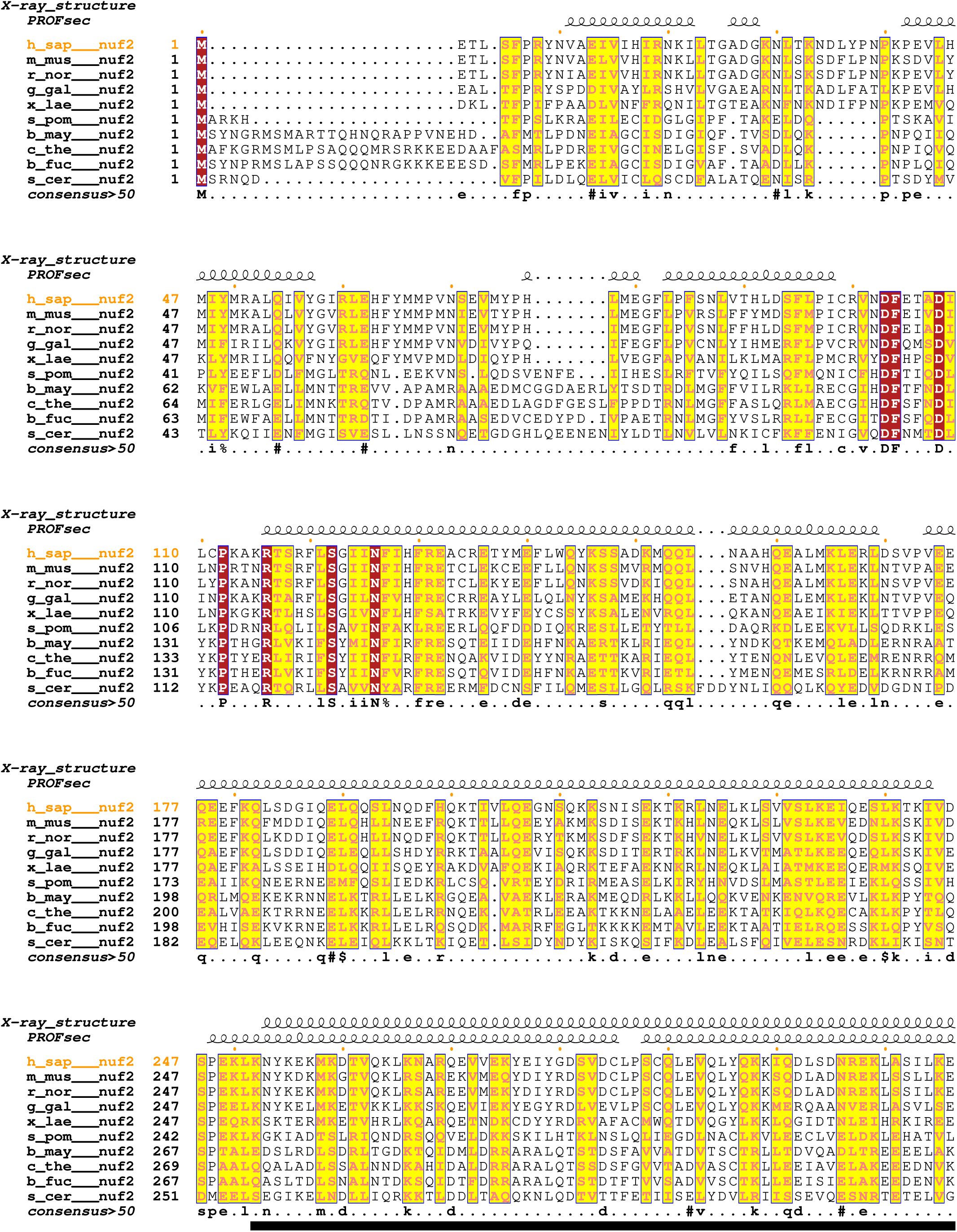
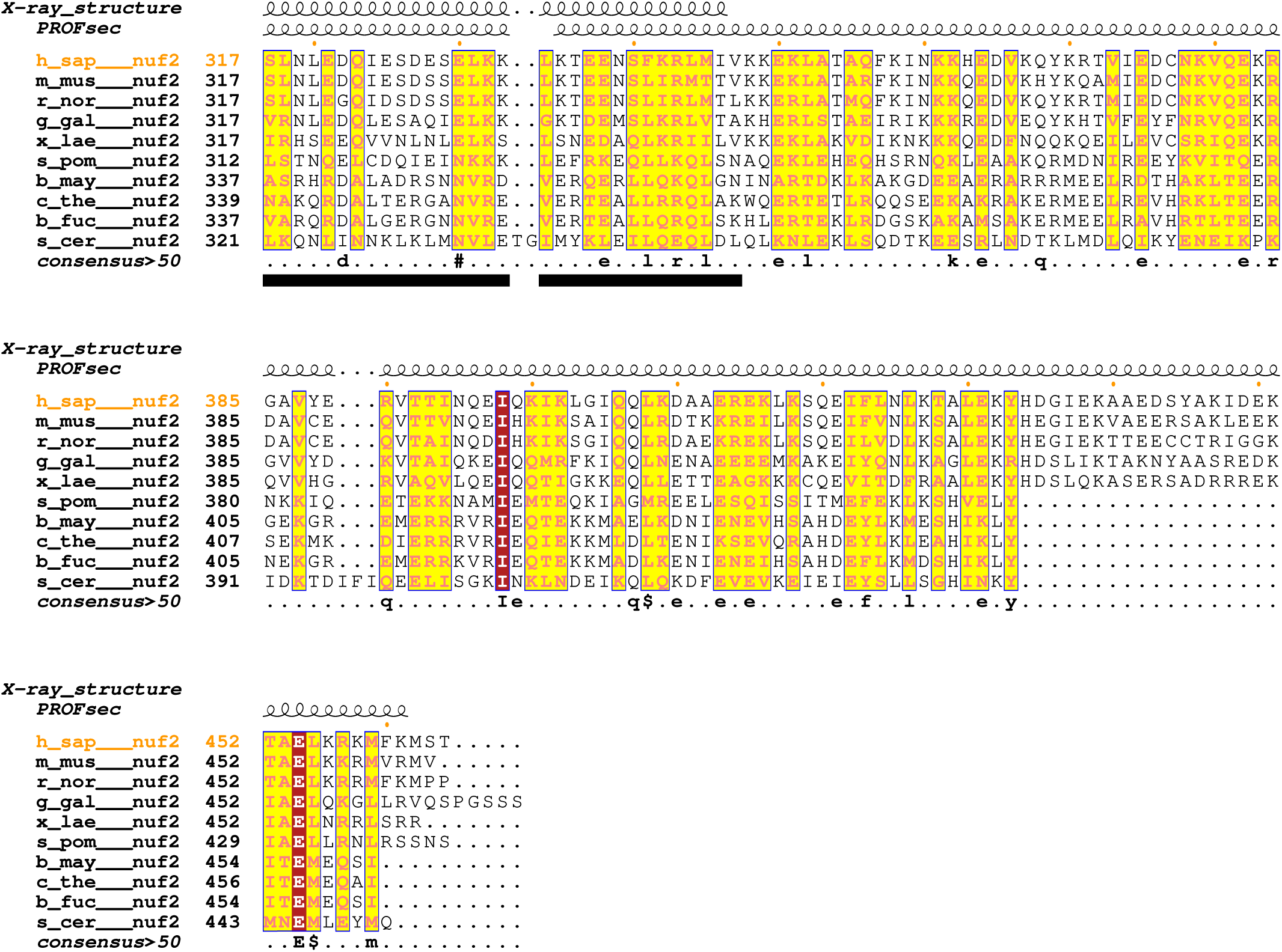
Nuf2 multiple-sequence alignment from metazoans and fungi. Sequences were retrieved from the UniProt (UniProt, 2021) database with the following accession identifiers: h_sap: *Homo sapiens*, Q9BZD4; m_mus: *Mus musculus*, Q99P69; r_nor: *Rattus norvegicus*, Q6AYL9; g_gal: *Gallus gallus*, Q76I90; x_lae: *Xenopus laevis*, Q6GQ71; s_pom: *Schizosaccharomyces pombe*, Q10173; b_may: *Bipolaris maydis*, M2TWT1; c_the: *Chaetomium thermophilum*, G0S7N6; b_fuc: *Botryotinia fuckeliana*, G2YM26; s_cer: *Saccharomyces cerevisiae*, P33895. Sequences were aligned with MAFFT (Katoh et al., 2002) and displayed with ESPript (Robert and Gouet, 2014). Above the sequences, secondary structure is shown as annotated by DSSP (Kabsch and Sander, 1983) from the Ndc80:Nuf2 loop crystal structure or as predicted with PROFsec (Rost, 2001), respectively. The black bars below the sequences are residues that were modeled in the Ndc80:Nuf2 loop crystal structure here.

**Data S3.**
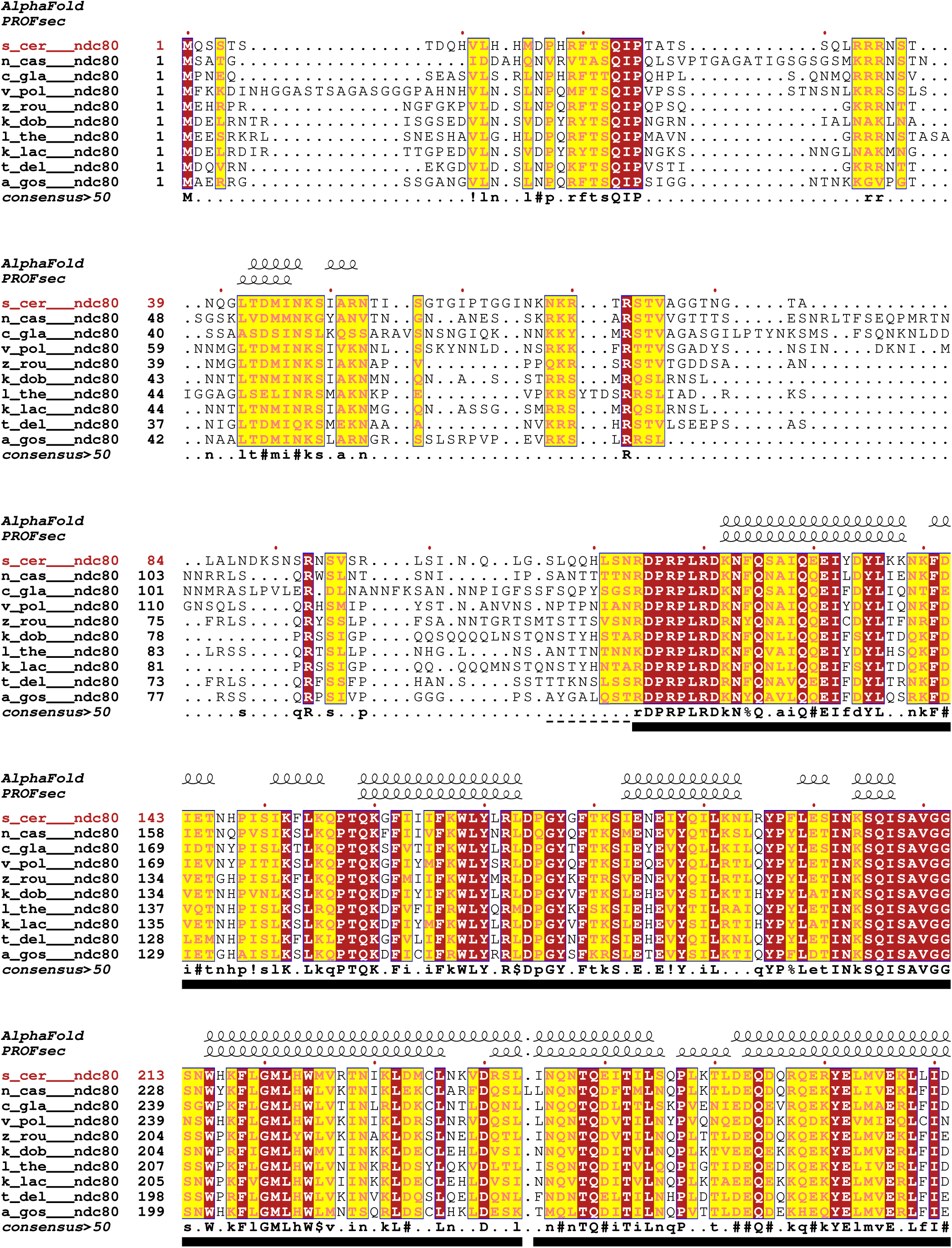
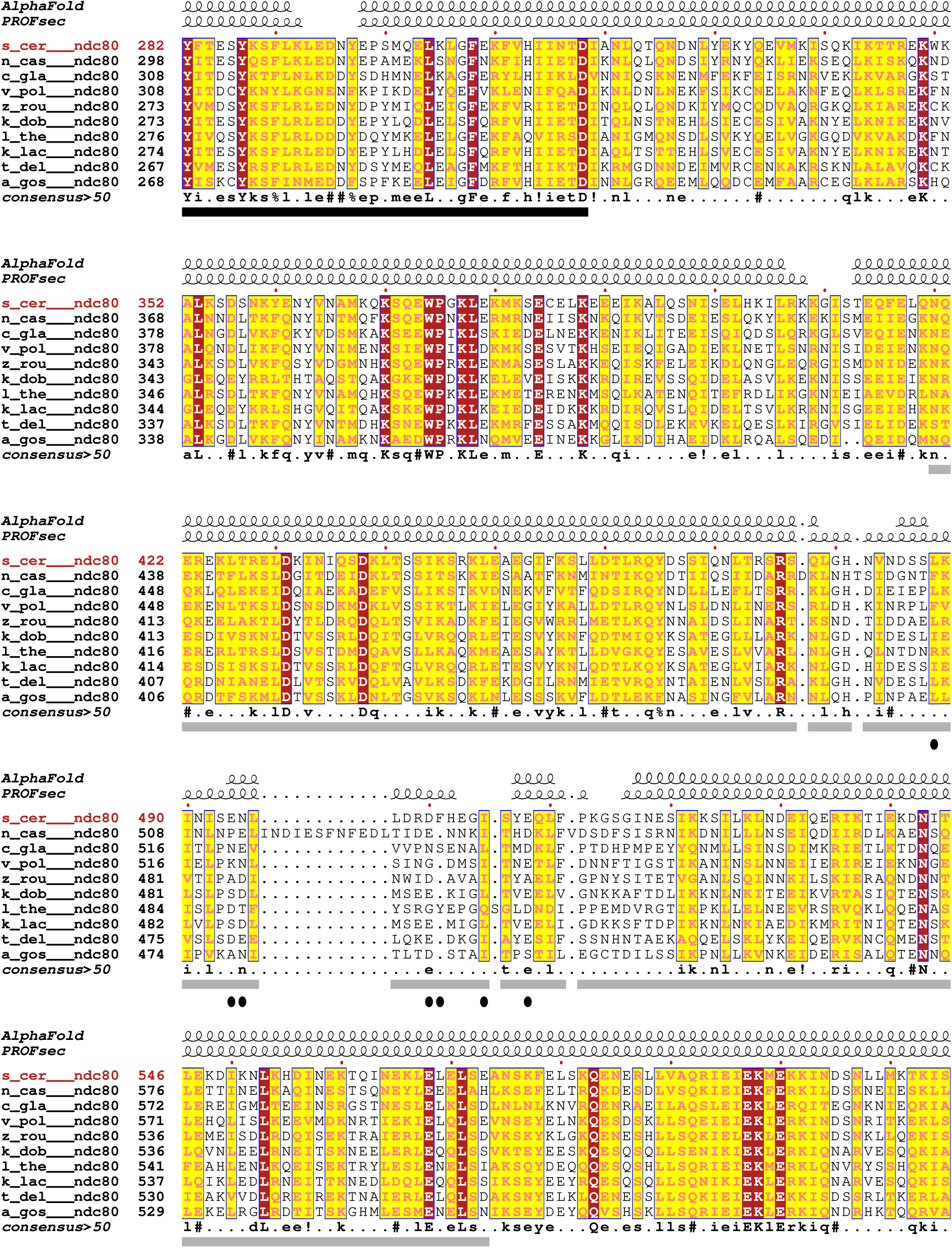
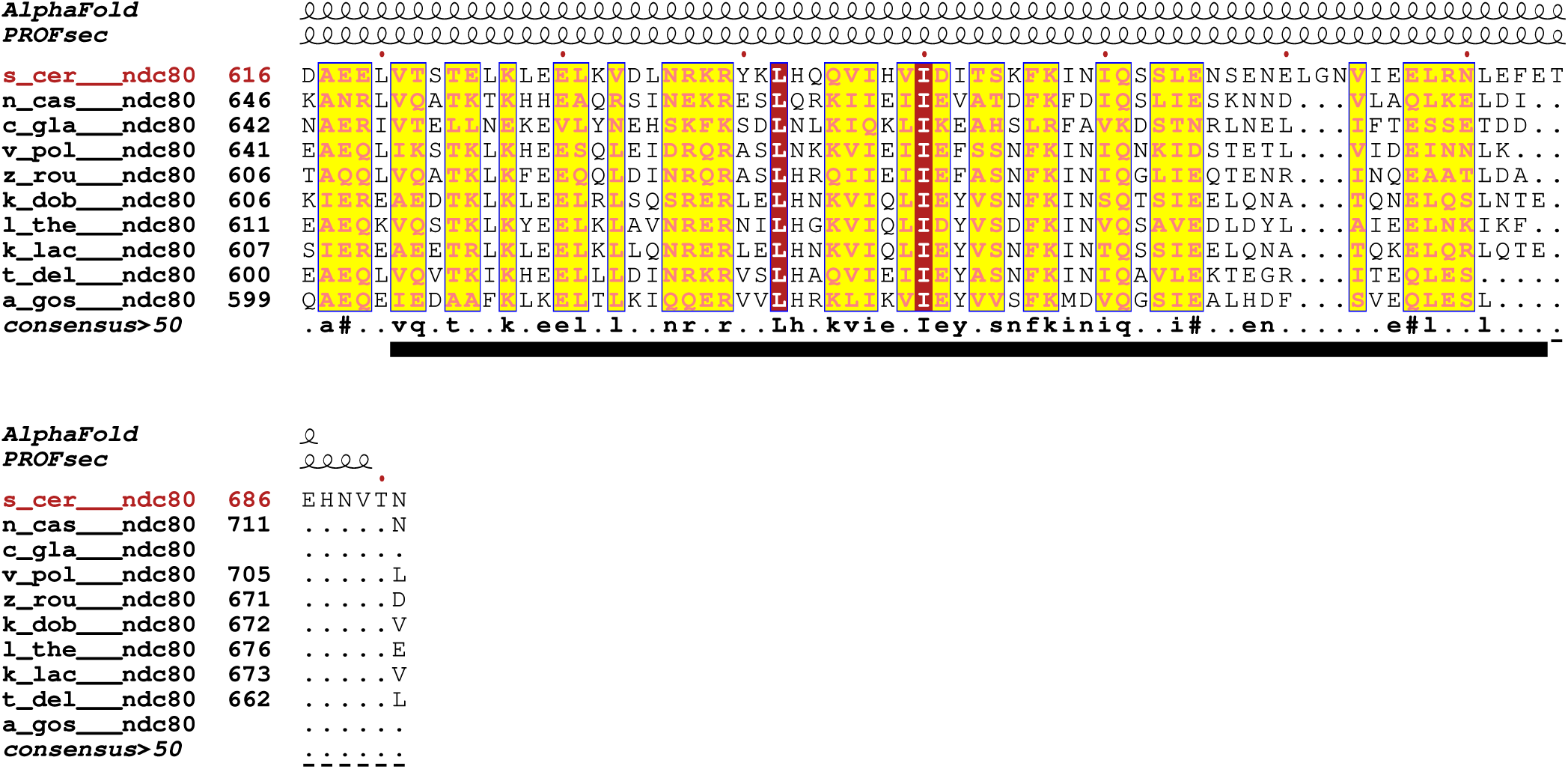
Ndc80 multiple-sequence alignment from Saccharomycetaceae budding yeast species. Sequences were retrieved from the UniProt (UniProt, 2021) database with the following accession identifiers: s_cer: *Saccharomyces cerevisiae*, P40460; n_cas: *Naumovozyma castellii*, G0VHN1; c_gla: *Candida glabrata*, Q6FW62; v_pol: *Vanderwaltozyma polyspora*, A7TQ22; z_rou: *Zygosaccharomyces rouxii*, A0A1Q3A623; k_dob: *Kluyveromyces dobzhanskii*, A0A0A8L398; l_the: *Lachancea thermotolerans*, C5DH42; k_lac: *Kluyveromyces lactis*, Q6CNF3; t_del: *Torulaspora delbrueckii*, G8ZP11; a_gos: *Eremothecium gossypii*, Q753N1. Sequences were aligned with MAFFT (Katoh et al., 2002) and displayed with ESPript (Robert and Gouet, 2014). Above the sequences, secondary structure is shown as annotated by DSSP (Kabsch and Sander, 1983) from the predicted AlphaFold model (Jumper et al., 2021) or as predicted with PROFsec (Rost, 2001), respectively. The gray bar below the sequences indicates the Ndc80 residues for which the crystal structure of the corresponding segment of the human loop structure was determined here. Black bars below the sequences are residues that were modeled in the Ndc80:Dam1-Nuf2 crystal structure here, black dashed lines are residues that were included in the construct but not modeled. Residues mutated to alanine in the *ndc80-7A* (Maure et al., 2011) mutant are indicated by black ovals (488, 494, 495, 500, 501, 505, and 508).

**Data S4.**
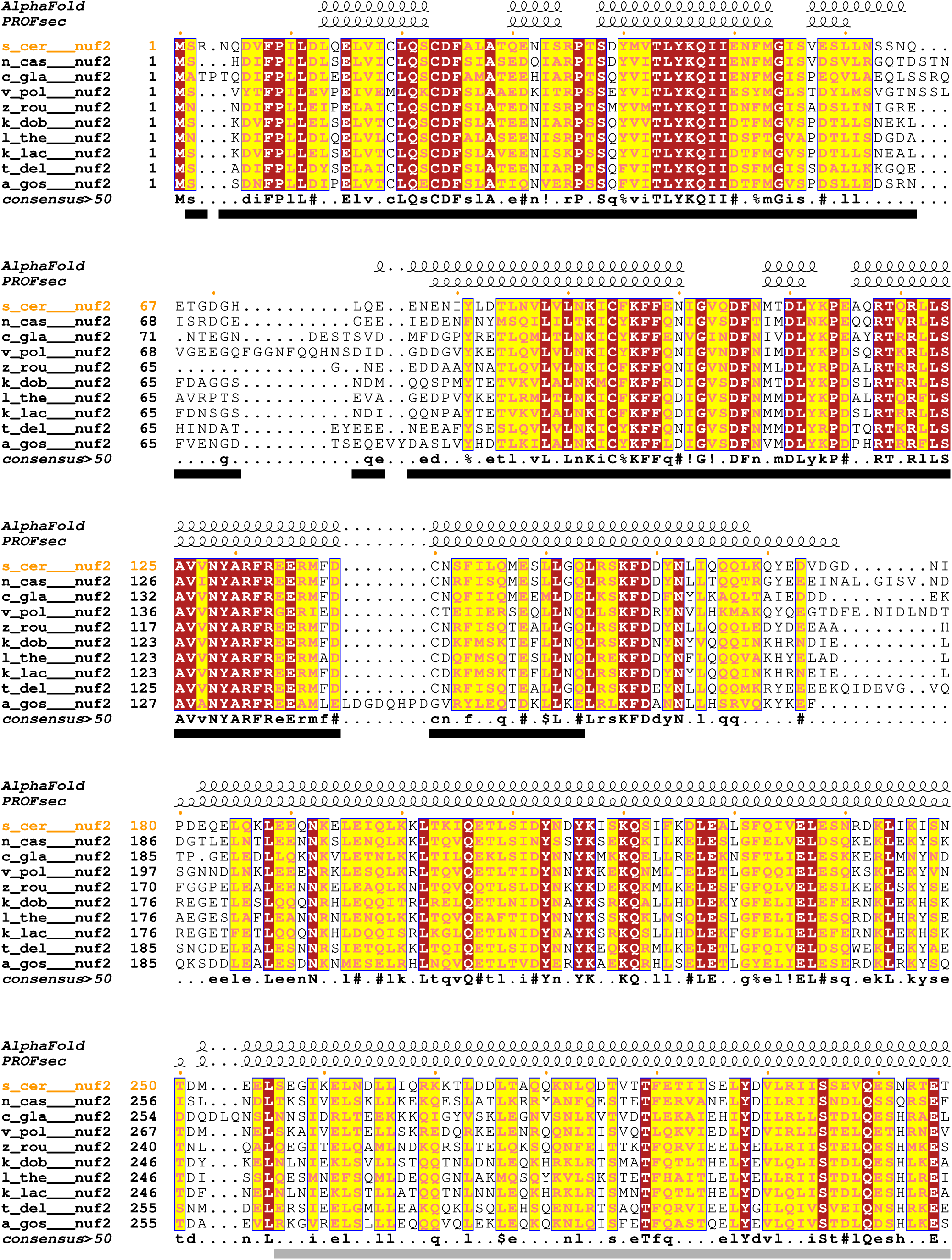
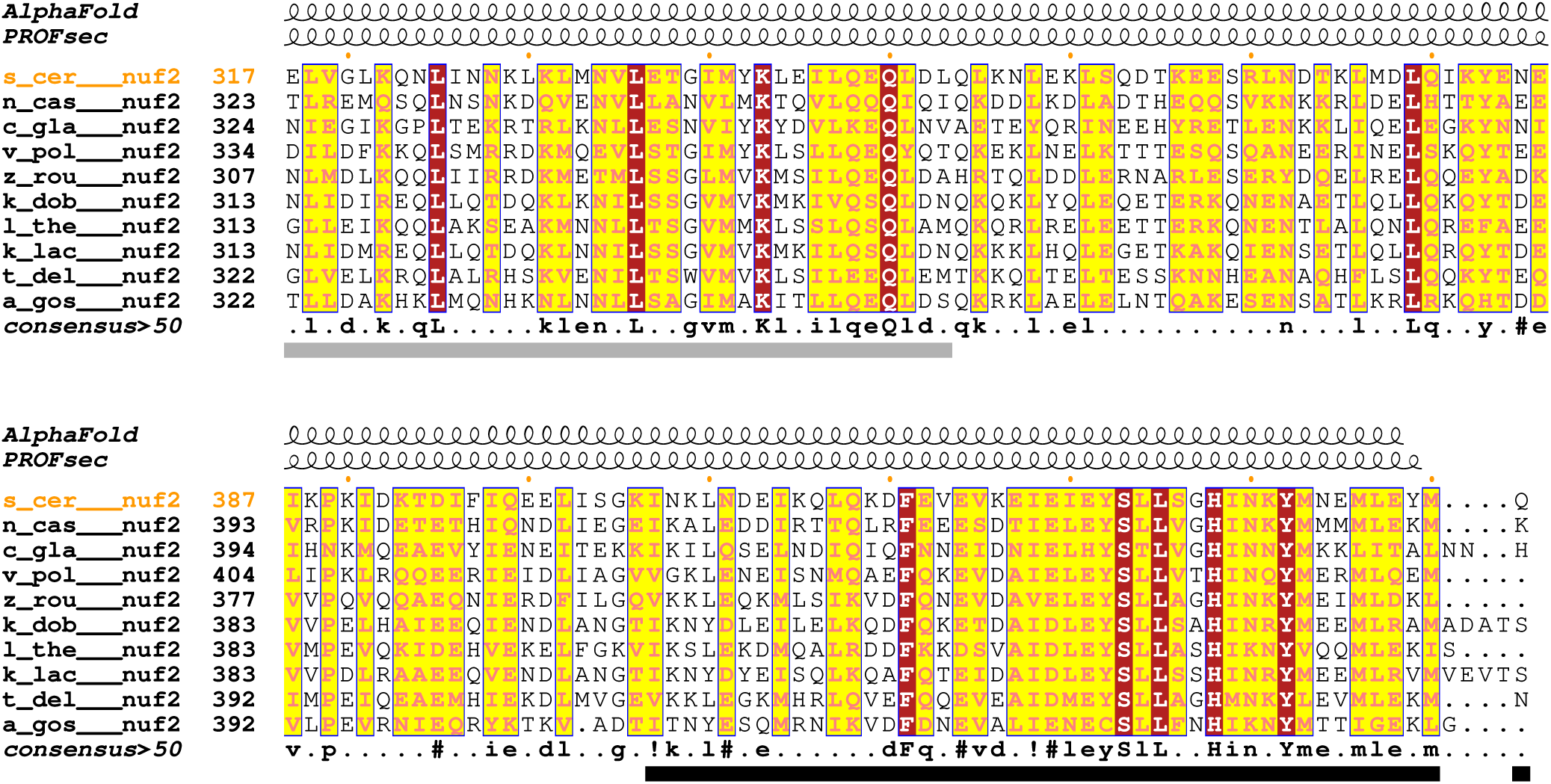
Nuf2 multiple-sequence alignment from Saccharomycetaceae budding yeast species. Sequences were retrieved from the GenBank (Benson et al., 2013) and UniProt (UniProt, 2021) database with the following accession identifiers: s_cer: *Saccharomyces cerevisiae*, P33895; n_cas: *Naumovozyma castellii*, G0VCX7; c_gla: *Candida glabrata*, Q6FNH8; v_pol: *Vanderwaltozyma polyspora*, A7TGX5; z_rou: *Zygosaccharomyces rouxii*, A0A1Q3A9H1; k_dob: *Kluyveromyces dobzhanskii*, CCBQ010000047.1, where the correct open reading frame was obtained with AUGUSTUS (Stanke and Morgenstern, 2005); l_the: *Lachancea thermotolerans*, C5DEF0; k_lac: *Kluyveromyces lactis*, Q6CJ06; t_del: *Torulaspora delbrueckii*, G8ZTW3; a_gos: *Eremothecium gossypii*, Q757M3. Sequences were aligned with MAFFT (Katoh et al., 2002) and displayed with ESPript (Robert and Gouet, 2014). Above the sequences, secondary structure is shown as annotated by DSSP (Kabsch and Sander, 1983) from the predicted AlphaFold model (Jumper et al., 2021) or as predicted with PROFsec (Rost, 2001), respectively. The gray bar below the sequences indicates the Nuf2 residues for which the crystal structure of the corresponding segment of the human loop structure was determined here. Black bars below the sequences are residues that were modeled in the Ndc80:Dam1-Nuf2 crystal structure here.

**Data S5.**
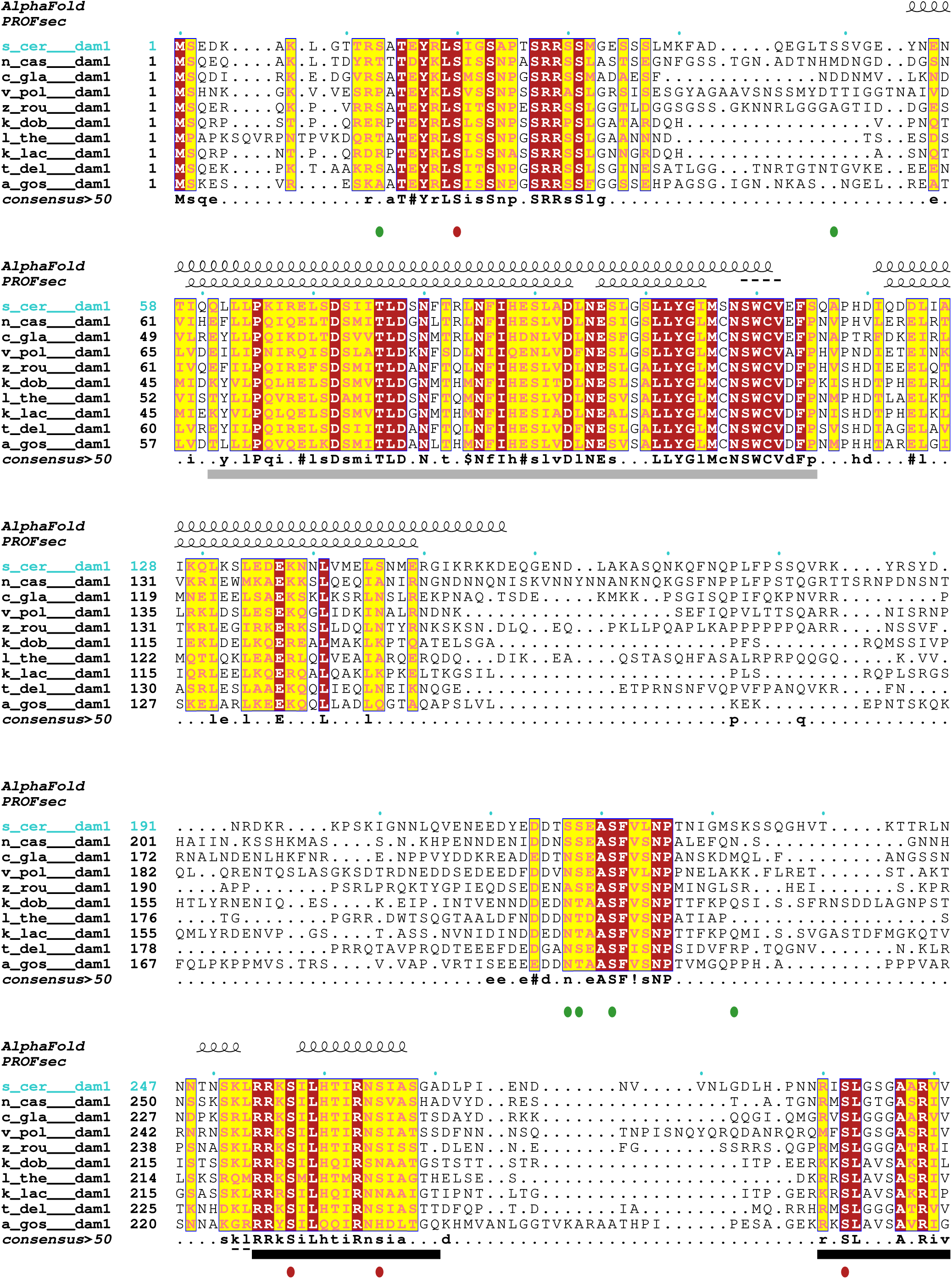
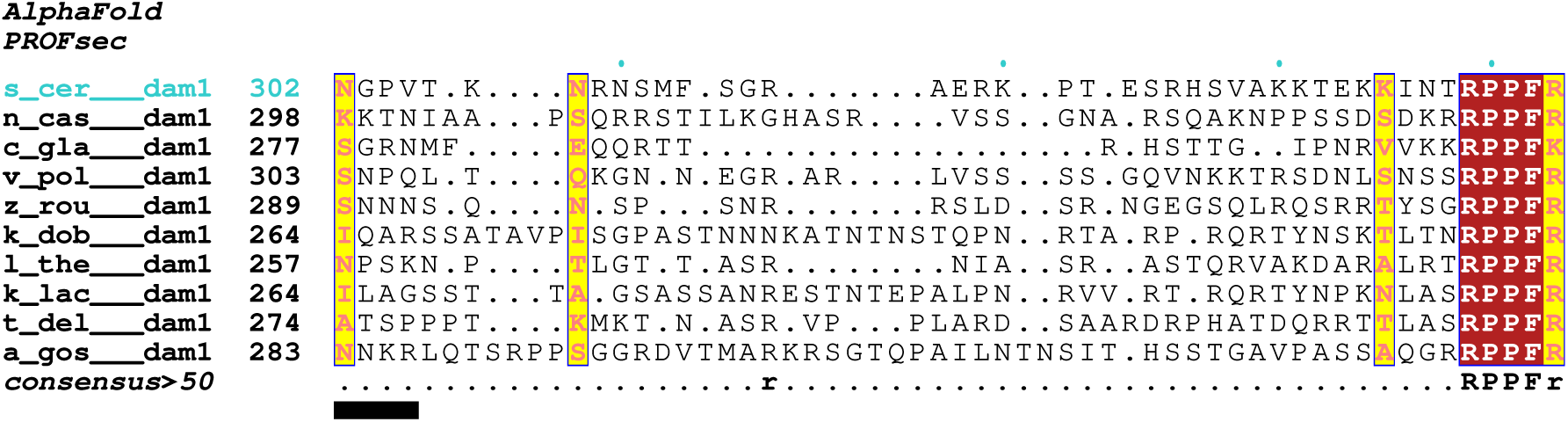
Dam1 multiple-sequence alignment from Saccharomycetaceae budding yeast species. Sequences were retrieved from the GenBank (Benson et al., 2013) and UniProt (UniProt, 2021) databases with the following accession identifiers: s_cer: *Saccharomyces cerevisiae*, NM_001181242_1; n_cas: *Naumovozyma castellii*, G0VF94; c_gla: *Candida glabrata*, XM_448982_1; v_pol: *Vanderwaltozyma polyspora*, XM_001647039_1; z_rou: *Zygosaccharomyces rouxii*, XM_002495884_1; k_dob: *Kluyveromyces dobzhanskii*, A0A0A8L2Q2; l_the: *Lachancea thermotolerans*, XM_002554750_1; k_lac: *Kluyveromyces lactis*, XM_452038_1; t_del: *Torulaspora delbrueckii*, G8ZX10; a_gos: *Eremothecium gossypii*, M_207923_1. Sequences were aligned with MAFFT (Katoh et al., 2002) and displayed with ESPript (Robert and Gouet, 2014). Above the sequences, secondary structure is shown as annotated by DSSP (Kabsch and Sander, 1983) from the predicted AlphaFold model (Jumper et al., 2021) or as predicted with PROFsec (Rost, 2001), respectively. The gray bar below the sequences indicates corresponding Dam1 residues that were modeled in the structure of the *Chaetomium thermophilum* DASH/Dam1c cryo-EM structure (PDB-ID 6CFZ) (Jenni and Harrison, 2018). Black bars below the sequences are residues that were modeled in the structure here, black dashed lines are residues that were included in the construct but not modeled. Phosphorylation sites are indicated by ovals: red, Ipl1 sites (Ser20, Ser257, Ser265, Ser292); green, Mps1 sites (Ser13, Ser49, Ser217, Ser218, Ser221, Ser232).

**Data S6.**
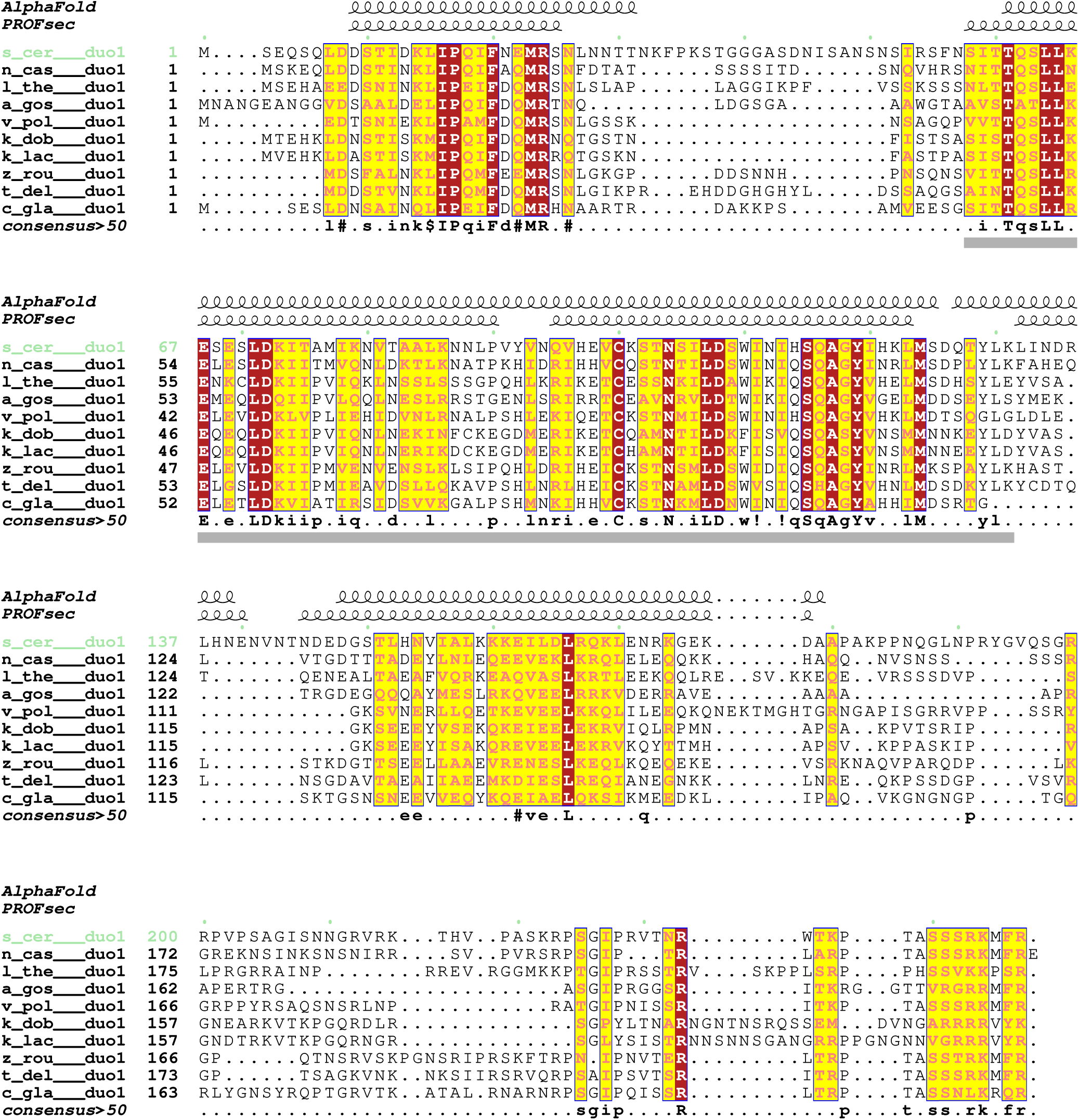
Duo1 multiple-sequence alignment from Saccharomycetaceae budding yeast species. Sequences were retrieved from the GenBank (Benson et al., 2013) and UniProt (UniProt, 2021) databases with the following accession identifiers: s_cer: *Saccharomyces cerevisiae*, NM_001180926_1; n_cas: *Naumovozyma castellii*, G0VEA9; c_gla: *Candida glabrata*, XM_446271_1; v_pol: *Vanderwaltozyma polyspora*, XM_001643060_1; z_rou: *Zygosaccharomyces rouxii*, XM_002495823_1; k_dob: *Kluyveromyces dobzhanskii*, A0A0A8L259; l_the: *Lachancea thermotolerans*, XM_002556121_1; k_lac: *Kluyveromyces lactis*, XM_452455_1; t_del: *Torulaspora delbrueckii*, G8ZXV6; a_gos: *Eremothecium gossypii*, NM_208057_1. Above the sequences, secondary structure is shown as annotated by DSSP (Kabsch and Sander, 1983) from the predicted AlphaFold model (Jumper et al., 2021) or as predicted with PROFsec (Rost, 2001), respectively. The gray bar below the sequences indicates corresponding Duo1 residues that were modeled in the structure of the *Chaetomium thermophilum* DASH/Dam1c cryo-EM structure (PDB-ID 6CFZ) (Jenni and Harrison, 2018).

## Notes

### Competing Interest Statement

The authors have declared no competing interest.

